# Machine Learning-Based Identification of B4GALNT1 as a Key Player in Hepatocellular Carcinoma: A Comprehensive Bioinformatics and Structural Analysis

**DOI:** 10.1101/2024.01.29.577885

**Authors:** Rohit Kumar Verma, Kiran Bharat Lokhande, Prashant Kumar Srivastava, Ashutosh Singh

**Author notes:** **Corresponding Author Details- Dr. Prashant Shrivastava** National Heart and Lung Institute, Imperial College London, London, United Kingdom.; **Dr. Ashutosh Singh** Associate Professor, Translational Bioinformatics and Computational Genomics Laboratory, Department of Life Sciences, School of Natural Sciences (SONS), Shiv Nadar Institution of Eminence (SNIoE), Delhi NCR, India.

## Abstract

Liver hepatocellular carcinoma (LIHC) is one of the most frequent types of malignant cancer in the globe. The identification of new biomarkers for the LIHC is critical. We used TCGA-LIHC gene expression datasets for this study. Several feature selection methods were used to find the top gene signatures that distinguish LIHC cancer from normal samples. Eleven machine learning algorithms were used on these selected characteristics, and model performance evaluation revealed that Naive Bayes Classifiers (AUC = 0.965) performs the best for a selection of 55 protein coding genes. Among 55 protein coding genes we found B4GALNT1 (Beta-1,4-N-acetyl-galactosaminyltransferase 1) which is differentially regulated in LIHC. With several evidence B4GALNT1 plays crucial role in tumorigenesis in many cancers, therefore we conducted systematic bioinformatics approach with mutational and structural analysis of B4GALNT1 in LIHC. Moreover, survival analysis, immune cell infiltration, most significant associated methylated CpG probe and access the accuracy of B4GALNT1 conducted to find the potential role of B4GALNT1. The results suggested that B4GALNT1 was significantly expressed in most cancers including LIHC. Finally, 16 missense mutations identified through cBioportal, Cosmic Database, and Human Variant Database, among which 6 mutations (P64Q, S131F, A311S, R340Q, D478H, and P507Q) found to be deleterious when analysed by *in-silico* prediction algorithms such as SIFT, PolyPhen2, I Mutent2 and CADD in LIHC. Molecular Dynamics simulation analysis was performed to understand the atomic details of the structure and functional changes. Results from this study suggest the impact of these missense variants on the structure of the B4GALNT1 protein and its pathogenic relevance. Our study demonstrated that B4GALNT1 may be evaluated as a novel target for liver cancer therapy because it has been found to be overexpressed in Liver and correlates with a poor prognosis.

## Introduction

Cancer stands as one of the most fatal primary contributors to global mortality. Among which breast, lung, colorectum, prostate and stomach are the top 5 leading causes and deaths in cancer were observed globally [1]. Liver hepatocellular carcinoma (LIHC) is the most frequent primary malignant tumour of the liver. The global prevalence of LIHC is rising. LIHC is common in people with cirrhosis and chronic liver disease [2]. LIHC ranks as the sixth most prevalent cancer worldwide and holds the position of the third primary factor leading to cancer-related deaths. According to Globacan report, the expected number of deaths in LIHC is around 830,180 individuals in 2020, accounting for around one-third of all cancer deaths [3]. Globally, the number of LIHC deaths is expected to rise to 1.24 million by 2030. The main cause of liver cancer deaths is that it often diagnosed at late stages and early detection of liver cancer might reduce the curve of death. LIHC is the most frequent kind of liver cancer in East and Southeast Asia and Sub-Saharan Africa, accounting for more than 80% of all cases and mortality. The high frequency of chronic hepatitis B virus (HBV) infection is partly responsible for the high incidence of LIHC in these areas. HBV is expected to be responsible for roughly 61% of all LIHC cases worldwide in 2020 [4]. Abdominal discomfort and weight loss are the most prevalent signs of LIHC. Imaging investigations, such as computed tomography (CT) or magnetic resonance imaging (MRI), are commonly used to diagnose LIHC (MRI) [5]. We tried to provide an insight to B4GALNT1 to carry out bioinformatics analysis for LIHC from The Cancer Genome Atlas (TCGA) [6]. Recent study has shown that in liver cancer, B4GALNT1 promotes carcinogenesis by regulating epithelial– mesenchymal transition (EMT) [7] indicating B4GALNT1 might be a potential biomarker for liver cancer. Data mining and techniques has widened the horizon for medical research by accessing the publicly available clinical samples (TCGA) and access the patient’s risk and to understand the role of particular genes in cancer.

β-1,4-N-acetyl-galactosaminyltransferase 1 is a transferase protein involved in glycosphingolipid biosynthesis that is required for the cell’s normal physiological functioning. In many different types of cancer, abnormal expression of particular GSLs and associated enzymes are related to tumour initiation and malignant transformation. B4GALNT1, also known for GM2/GD2 synthase involved in the formation of sialic acid-containing glycosphingolipids (GSLs) [8].

The change in expression or mutation B4GALNT1 has been associated with tumor progression in melanoma [9] by inducing of gangliosides GM2/GD2, clear cell renal cell carcinoma (ccRCC) [10], colorectal cancer [11], lung adenocarcinoma [12], oral squamous cell carcinoma [13], breast cancer [14] and cervical cancer [15]. Apart from cancer B4GALNT1 has been associated with Hereditary Spastic Paraplegia [16], Parkinson’s disease [17] and Axonal Charcot-Marie-Tooth Disease [18]. The aforementioned findings indicate the significance of B4GALNT1 as a crucial regulator in the progression of cancer. The aberrant expression of B4GALNT1 has the potential to impact the prognosis of individuals with cancer. Consequently, there is a need for a thorough and structural analysis of B4GALNT1 in LIHC.

In the present study, a machine learning and systems biology method were employed to discover LIHC-associated genes that may serve as prognostic indicators, and we conducted comprehensive and structural analysis of B4GALNT1 in LIHC. The TCGA, TIMER2 and UALCAN has been used to find the expression level of B4GALNT1 in different types of cancers. We also explored the relationship between the B4GALNT1 and the immune cell infiltration. Structural analysis conducted in order to study the impact of mutation in the structure through molecular dynamic simulation in LIHC. We identified 16 missense mutations were identified through cBioportal [19], Cosmic Database [20], and Human Variant Database (HuVarBase) [21], among which 6 mutations (i.e., P64Q, S131F, A311S, R340Q, D478H, and P507Q) found to be deleterious using *in-silico* prediction algorithms such as SIFT, PolyPhen2, I Mutent2 and CADD.

## Materials and Methods

**Figure 1** provides a concise overview and description of the study’s workflow, outlining the process from obtaining LIHC datasets from TCGA to conducting a thorough structural analysis of B4GALNT1.

**Figure 1.**
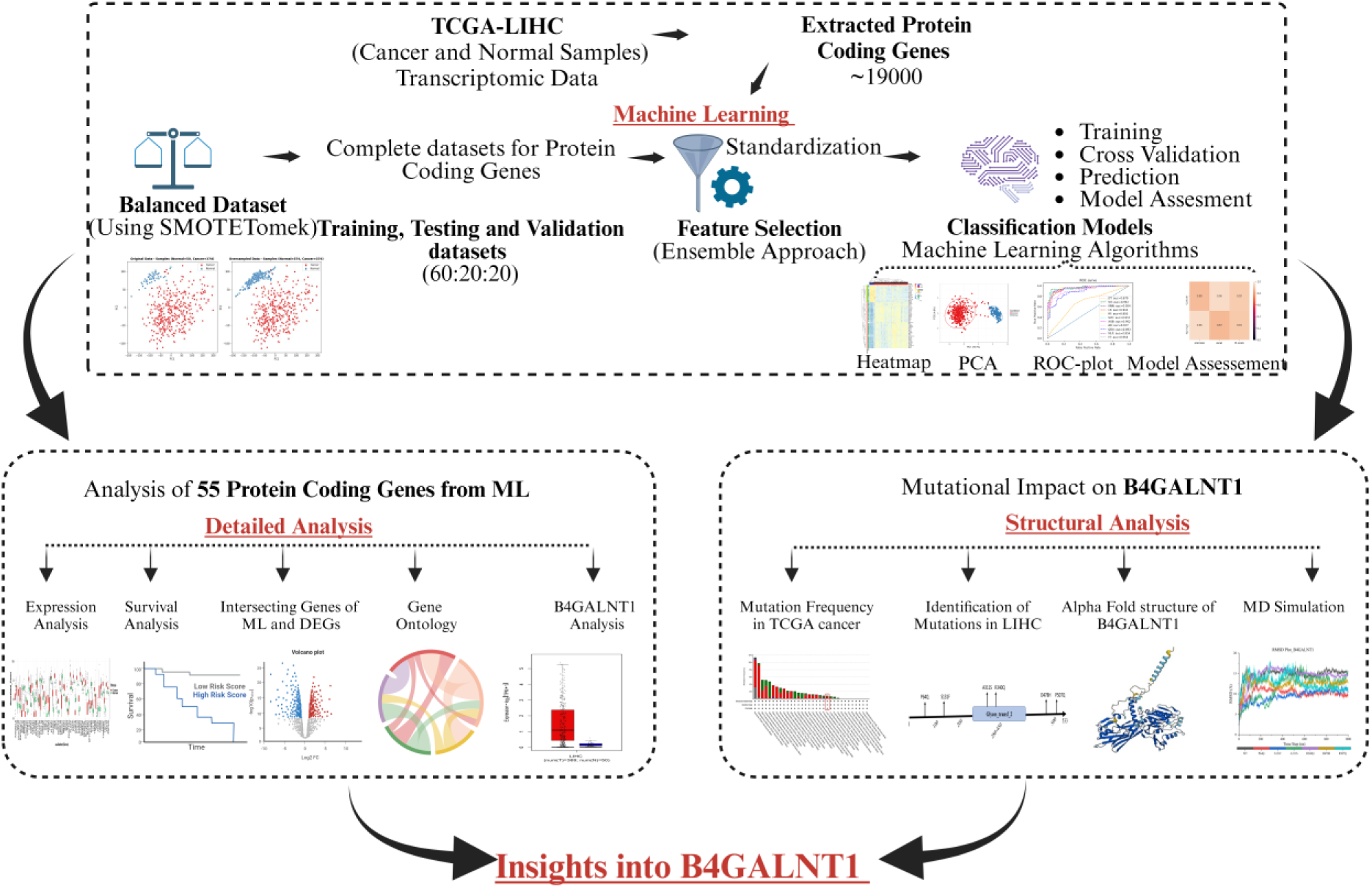
Represent a schematic illustration of LIHC’s bioinformatics analysis. In this work, TCGA-LIHC transcriptome datasets (Normal=50, Cancer=374) were used. From the LIHC transcriptome dataset (19963), only protein coding genes were retrieved. The SMOTETomek method is utilised to over sample the minority class (Normal=374, Cancer=374). Furthermore, the datasets have been separated into 60:20:20 ratios for training, testing, and validation. For genes that distinguish cancer from normal samples, a different feature selection strategy was used. A machine learning technique was used for classification and performance measurement. Differential gene expression analysis as well as other bioinformatics analyses were carried out. In addition, we employed B4GALNT1 from 55 protein-coding genes for compressive and structural analysis.

### Transcriptome dataset (LIHC) download and pre-processing

In this study, we have accessed publicly available datasets for LIHC from TCGA and downloaded the transcriptome dataset (Count dataset). The dataset comprising 424 samples (Normal= 50 and Cancer= 374). Further, we retrieved the dataset with protein coding genes (Protein coding= 19963). Finally, the protein coding genes were merged to samples with mapped to their corresponding Ensembl IDs. This selection results in a protein-encoding gene matrix of 19963. The dataset was normalized using cpm normalization method for machine learning analysis in R/RStudio using edgeR [22]. After pre-processing, the combined dataset with solely protein coding genes yields an initial dataset with 424 samples and 19963 features, referred to as the complete dataset.

### Machine Learning

Since normal samples were very few (Normal= 50 and Cancer= 374) as compared to cancer samples therefore we employed SMOTETomek [23] to oversample the minority class using in-house Python script. SMOTETomek is a hybrid approach in machine learning for uneven class distribution. SMOTETomek uses two different algorithms SMOTE (Synthetic Minority Over-sampling Technique) and Tomek links. SMOTE oversample the minority class and Tomek first identify and then removes the potential noisy instances from the dataset using Tomek links. A balanced dataset obtained after applying SMOTETomek algorithm (Normal= 374 and Cancer= 374).

### Feature Selection Methods

One of the major issues in this study is identifying the crucial set of characteristics for machine learning models. Only a few gene groups are relevant to detect specific disease or biological activity from thousands of genes that can be detected from next generation sequencing methods. Feature selection is the act of automatically selecting those features in the data that contribute the most to the prediction variable. It is an important stage in machine learning since it helps you to limit the amount of data that the model must learn, which speeds up training and improves performance. It also enables to discover and delete characteristics that are irrelevant to the job at hand, which might enhance the model’s interpretability. The detailed feature selection method has been provided in Supplementary methodology section.

### Machine Learning Models

After pre-processing and employing feature selection methods we employed several machines learning algorithms, including K-Nearest Neighbours (KNN), Random Forest (RF), Support Vector Machine (SVM), AdaBoost (AB), Quadratic Discriminant Analysis (QDA), Decision Tree (DT), XGBoost (XGB), ExtraTrees (ET), Gaussian Naive Bayes (GNB), Logistic Regression (LR), and Multi-layer Perceptron (MLP) were used in this study to develop binary classification models. We fine-tuned the model on training data by using several probability thresholds such as 1.0, 0.9, 0.8, 0.7, 0.6, 0.5, 0.4, 0.3, 0.2, and 0.1, and we discovered that 0.5 works well for all models. The performance of the approaches was examined in order to select the best performing algorithm, which was then evaluated further. We performed the modelling procedure ten times for each technique and utilised a confusion matrix (CM) to present the classification results. **Figure 2** represents the overall workflow of Machine learning and the implementation of feature selection methods [24, 25].

**Figure 2.**
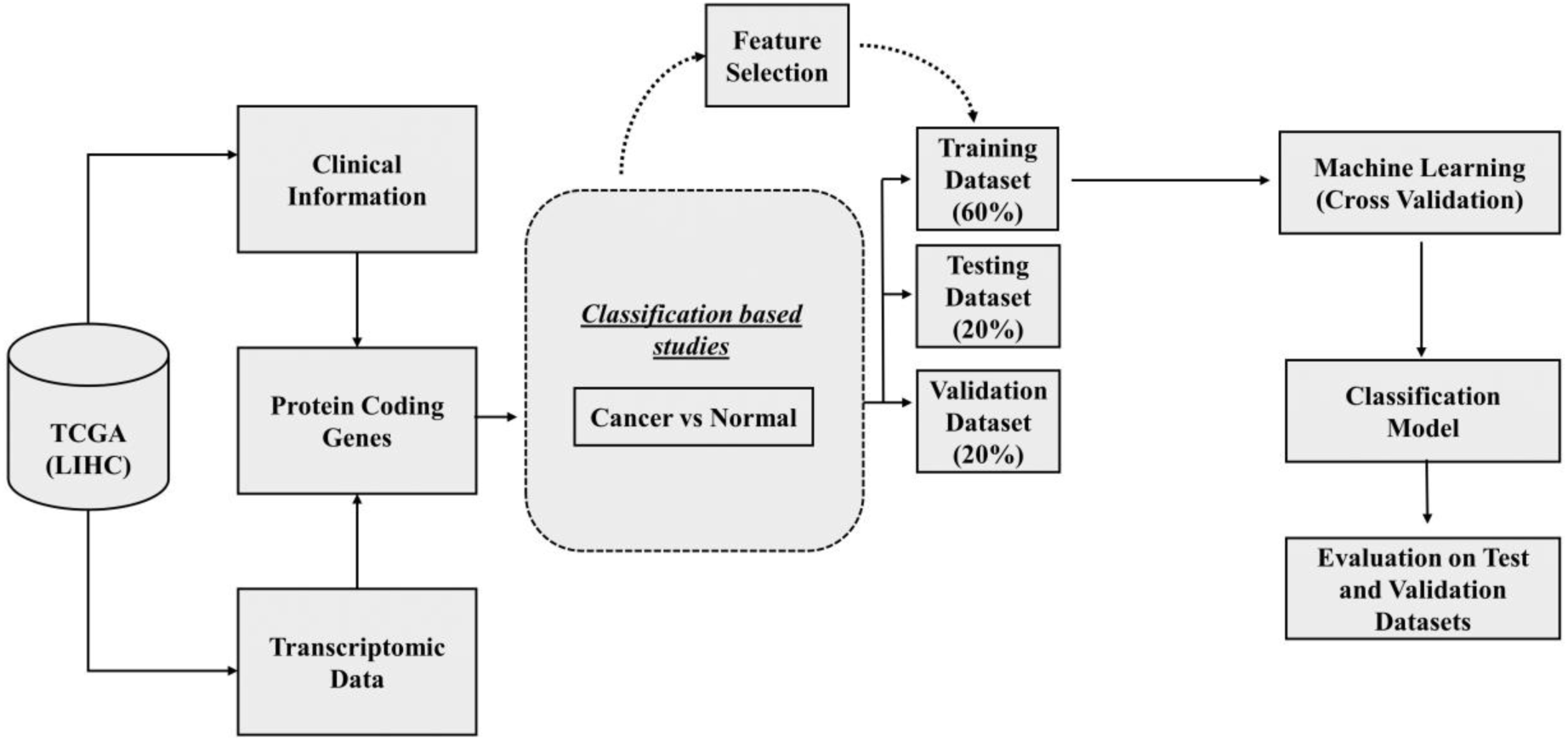
Represent the workflow of different feature selection methods and Machine Learning techniques.

### Training classification model and cross validation

The dataset was divided in a 60:20:20 ratios, with 60% of the data being training data and the remaining 20, 20% being test and validation datasets. Classification Models are built onto each and every feature selection technique and 5-Fold Cross-Validation is performed on training data set where model trains on 4 folds and predicts on the 5^th^ fold and this iteration runs on every fold and is averaged out. Internal validation is done on the test data. Results are noted to keep track of models which give promising results. In the fivefold cross-validation approach, the total training data is divided into five equal folds. During each iteration, four folds are employed for training, while the remaining fifth fold is designated for testing. This process is repeated five times, allowing each fold to take on the role of testing data at least once. Further, the performance measure was measured for test and validation datasets.

### Differential gene expression analysis (DGE)

Differential gene expression was evaluated by DESeq2 package in R/biocoductor [30] LIHC samples and normal tissues. The statistical threshold to determine DEGs using with a cut-off of |log2-fold change| > 1 and pvalue < 0.05. Further, volcano plot was generated to visualize the DEGs between LIHC and normal samples [26].

### Bioinformatics analysis

Several bioinformatics analyses were performed among which Kaplan-Meier survival analysis was done for all the 55-protein coding from ML-identified genes using KM plotter, an open-source web-based tool [27]. For survival analysis, patient samples in LIHC were categorized into high and low mRNA expression groups for each gene. Subsequently, Cox regression analysis was utilized to assess the association between prognostic genes and the outcomes of patient survival. A p-value of 0.05 was considered statistically significant for survival analysis and Cox regression analysis. For DEGs analysis and other analyses, R was used for all statistical tests, with a significance threshold of pvalue < 0.05. To determine the significance of differences between LIHC and normal samples, a paired t-test was used.

### Association of B4GALNT1 with pan-cancer and disease prognosis

The differential expression of B4GALNT1 was observed in different tumor and normal samples using CancerLivER (https://webs.iiitd.edu.in/raghava/cancerliver) [28], TIMER2 (http://cistrome.Shinyapps.io/timer/) [29], UALCAN database (http://ualcan.path.uab.edu) [30]. We also used UALCAN portal for examining the relative expression of B4GALNT1 in LIHC and normal tissues, as well as in other sub-groups such as cancer stages, tumour grades, age, gender, weight, historical subtypes, TP53 mutation status, promoter methylation, nodal metastasis status and race. Furthermore, we also used TNMplot (https://tnmplot.com/analysis/) [31] which provides differential gene expression of 22 cancers of tumor and normal samples. All of these online tools provide the analysis of differential expression of genes in different tumor and normal samples, which ultimately provides insight to understand B4GLANT1 gene expression in different types of cancers. To investigate the genes, which are positively and negatively correlated with B4GALNT1 expression in LIHC, we accessed LinkedOmics (https://linkedomics.org/login.php) which is an online application for analysing multi-omics and clinical data from the TCGA [32].

### Association of B4GALNT1 with prognosis in 33 different cancers

We conducted a comprehensive analysis of survival data obtained from UCSC Xena across 33 cancers. The assessment included overall survival, disease-free survival, disease-specific survival, and progression-free survival. The primary objective was to elucidate the correlation between the expression levels of the B4GALNT1 gene (categorized as high 50% and low 50%) and the progression of the disease in cancer patients across various cancer types. We plotted forest plot (R packages “survival” and “forestplot”) and survival plot (R packages “survival” and “survminer”) using Kaplan-Meier method [33] and log rank test were employed with significance p < 0.05 and we used month as survival time unit. We also used KM plotter, a web-tool for multivariate survival plot for gene expression of B4GALNT1 in LIHC [34]. The risk score and the lasso cox regression applied for 55 protein coding genes of each patient were calculated using sangerbox (http://vip.sangerbox.com/) [35].

### Gene Alteration analysis

The cBioportal (http://cbioportal.org) database was accessed for the mutation type, alteration frequency, and copy number alteration (CNA) from TCGA database [36]. Moreover, data for prognosis of B4GALNT1 in LIHC with or without genetic alteration was analysed using overall, disease-specific, disease-free, and progression-free data and obtained the plot with pvalues. Furthermore, we investigated different mutational site for B4GALNT1 in different resources such as Cosmic [37] and Human variant database [38], which is an open access web-based resource for accessing multidimensional genomics cancer datasets.

### Construction of PPI networks of B4GALNT1

We used GeneMANIA for finding association of B4GALNT1 and other proteins, genetic interactions, co-expression, co-localization and known and predicted pathways [39]. The PPI network of B4GALNT1 and the associated genes were constructed using STRING database [40]. The GO and pathway enrichment analysis were obtained using ShinyGO tool for 50 most interacted proteins from STRING database [41].

### Full atomistic molecular dynamic simulation

For the structural analysis B4GALNT1 model we used different *in-silico* based modelling, but we found the alpha-fold structure more suitable to study at atomic details for mutational aspect of structural impact. We downloaded the B4GALNT1 structure from uniport database and protein were subjected to residue mutation using Maestro [42].

We conducted a comprehensive molecular dynamics simulation for 1 μs (1000 ns) to assess the effects of mutations on the B4GALNT1 enzyme. This simulation encompassed the wild-type B4GALNT1 and its mutant variants, specifically P64Q, S131F, A311S, R340Q, D478H, and R507Q, utilizing the Desmond software [43]. We conducted a total of seven individual simulations, each spanning 1 μs, covering seven different structural configurations. These molecular dynamics simulations enable the calculation of forces and the tracking of the motion of amino acid atoms. It’s worth noting that Desmond offers enhanced capabilities, with a more comprehensive system for managing temperature, pressure, and volume, along with built-in functionalities for modelling protein dynamic behaviour [44]. The proteins were prepared within the Maestro program using Desmond’s system builder. In this process, they were inserted into an orthorhombic box filled with 1 Å spacing of water molecules. The water molecules were modelled using a three-point water model known as TIP3P. Additionally, a POPC lipid bilayer was introduced into the system, and periodic boundary conditions were applied. The placement of the POPC lipid bilayer was specifically on the N-terminal helix of the B4GALNT1, which encompassed residues Met1 to Arg28. The overall charge of the solvent system was balanced by introducing suitable counter ions at random within the solvated system. Energy minimization, a critical stage in MD, was performed using the steepest descent method. To mitigate edge effects in a finite system and enable the application of periodic boundary conditions, an orthorhombic box configuration was employed. This involved placing the atoms of the system to be simulated inside a space-filling box, which was then enveloped by replicated copies of itself. In this study, the OPLS_2005 force field, specifically designed for molecular dynamics simulations of proteins, was employed. Molecular dynamics simulations were executed with constraints set as "all-bonds" and utilized the MD integrator. Temperature control (300 K) was achieved using the Nose-Hoover chain thermostat, while pressure was regulated through the Martyna-Tobias-Klein barostat methods. The simulations were conducted within the NPT ensemble, ensuring constant particle number, pressure, and temperature. These parameters were carefully selected to accurately model the dynamic behaviour of proteins in the system.

Once the system reaches equilibrium, the MD simulation trajectories are recorded and analysed to investigate conformational alterations. Specifically, we examined the conformational changes in the C-alpha backbone of both the B4GALNT1 wild type and its mutant variants by comparing them to their initial conformations. This comparison was conducted using Root Mean Square Deviation (RMSD) and Root Mean Square Fluctuation (RMSF) plots. Additionally, we calculated changes in the side chains of the residues to provide a more comprehensive understanding of the structural dynamics.

## Results

### A precision gene panel for accurate cancer and normal sample discrimination in LIHC

Our major goal is to uncover possible biomarkers from protein coding genes that can classify between cancer and adjacent tissue samples. Following that, *in-silico* prediction models based on these characteristic markers were created utilising various machine learning methods. We extracted the protein coding genes from transcriptomic data and normalized the count data using limma edgeR package for further used for classification.

All 424 LIHC samples with 19963 protein coding gene features were used for the supervised machine learning classifiers to better classify cancer and normal tissue samples. Since the total sample size for LIHC dataset 424 tissue samples (50 normal and 374 LIHC) so, the datasets seem to be unbalanced. The cancer samples were almost 7 times larger than the normal samples. Therefore, we used SMOTETomek (Synthetic Minority Oversampling Technique) and Tomek link to balance the datasets (374 normal and 374 LIHC). The balanced dataset was split into training, testing and validation datasets in LIHC (in the ratio of 60:20:20). We employed rocc package to reduce the dimension of 19963 features in smaller subset and it subset the genes having AUC >= 0.90. This method identified 3149 feature genes which has AUC value >= 0.90. Furthermore, we employed variable importance from the model such as xgboost, random forest, support vector machine (linear), feature importance using Extra Tree classifier, boruta, univariate Spikeslab, fast correlation-based filter method (FCBF), Lasso Regression and Recursive Feature Elimination (RFE) to obtain the best gene features from 24,100,100,100,421,100,93,148,92 and 16. After that we have taken out the features that are common at least 50% of the total used feature selection method. Polling approach at the end provides 55 features from above mentioned datasets. Moreover, 55 genes obtained from the feature selection method were used as training dataset for eleven different classifiers i.e., AB, DT, NB, KNN, LR, RF, XGB, SVC, QDA, MLP and ET. For both the test and validation datasets, all the classifiers performed nearly well for 55 gene features. **Figure 3(A)** represents the separation of data of original dataset and the datasets after applying SMOTETomek. Further, PCA plot is generated depicting the separation of clusters (Cancer and normal samples) before and after SMOTETomek **Figure 3(B)**. Heatmap was used to visualize the 55 genes that could separate the LIHC, and normal samples based on the regulation of genes and ROC value **Figure 3(C)**. On the test datasets, we further examined the performance of all classifiers. The Naive Bayes classifier attained the best accuracy of 91.33% for 55 gene features for protein coding genes dataset, with an AUC of 0.96 **Figure 3 (D)** and **Supplementary Table S1**. Furthermore, for the validation dataset, the maximum accuracy was reached with AUCs of 0.98 and 0.95 for Logistic regression and Support vector classifier and accuracy was obtained 92.67% and 93.33%, respectively **Figure 3 (E)** and **Supplementary Table S2**. The DEGs were obtained using DESeq2 and volcano plot was used to visualize the genes that were intersecting with genes (Upregulated and Downregulated genes) obtained after DEGs analysis and the 55 ML-genes **Figure 3 (F).** Out of 55 genes 3 genes showed no regulation when compared with foldchange 2. The box plot is shown in the **Figure 3(G)** with regulation for cancer and normal samples.

**Figure 3:**
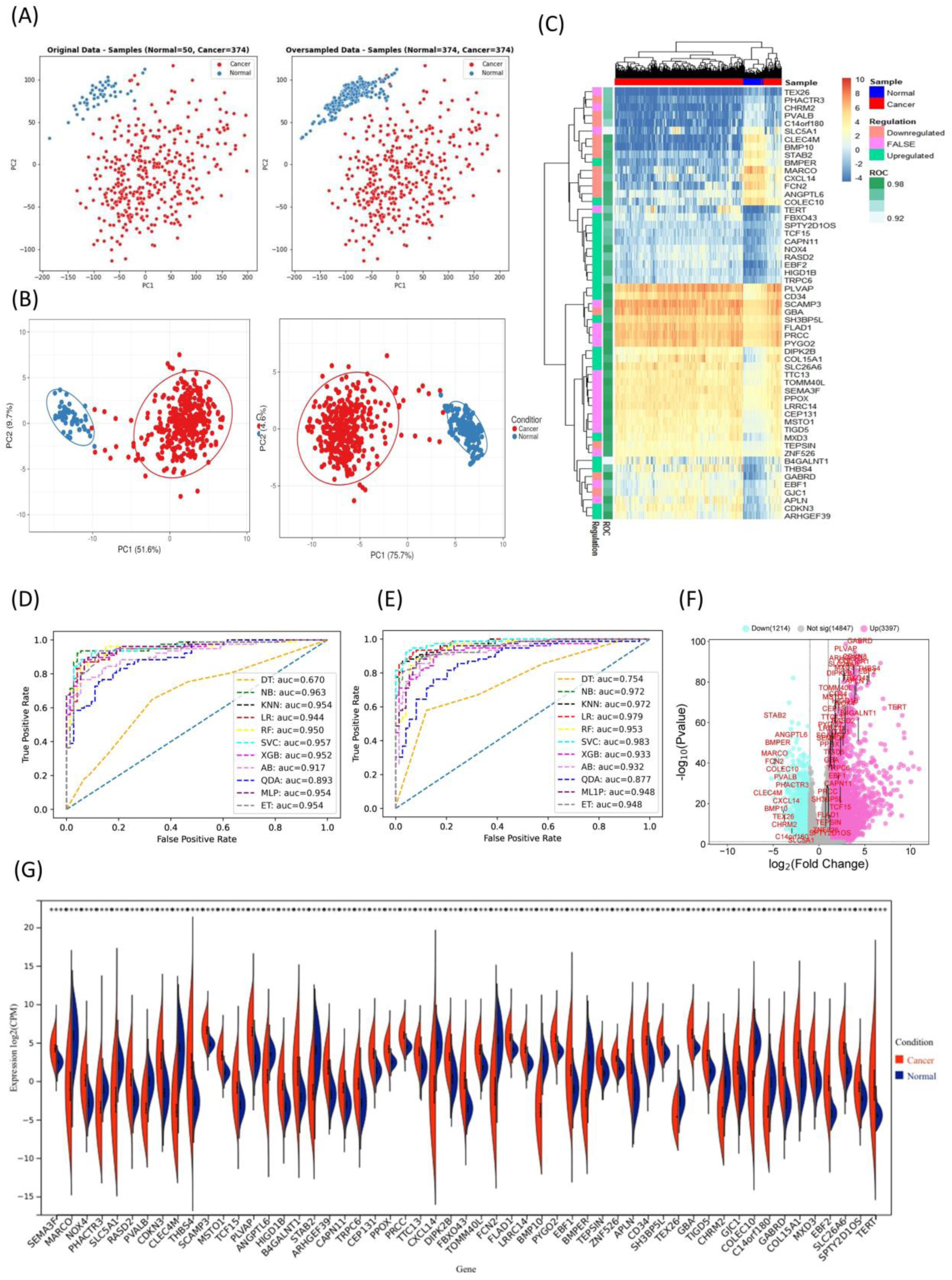
PCA plot depicting the two different clusters of cancer and normal samples (A-B) before and after applying SMOTETomek. (C) Heatmap showing the differential expression pattern in cancer and normal with regulation and ROC value. (C-D) ROC curve of the 55 PC genes for test and validation datasets. (E) Volcano plot for patients with LIHC compared with the normal with a threshold of log2FC > 1 in purple are upregulated and log2FC < 1 in cyan is downregulated. The ML obtained genes are represented in the volcano plot. (G) Complete differential expression of ML obtained genes are represented in violin plot.

Further to understand the significance of each feature from the ML-generated features in LIHC samples with their log2 Fold change, AUC, Pvalue, padj and Kaplan-Meier Log Rank P-value is provided in **Supplementary Table S3**.

### Protein-protein interaction and survival analysis

To study the physical interactions among 55 protein coding genes. The string online webserver tool was accessed with minimum required interaction score with confidence of 0.150 and we observed number of edges as 132 with an average node degree 4.8 **Figure 4(A)**. Using lasso-penalized cox regression the 55 genes were reduced to 13 genes with λ=0.04 **Figure 4(B)**. Out of 55 protein coding genes 20 genes were involved in KEGG pathways phagosome, mineral absorption, glycosphingolipid biosynthesis, hepatocellular carcinoma, pathways in cancer and other pathways shown in **Figure 4(C)**. For individual model, risk score was calculated for all LIHC patients based on expression value of 13 genes and based on the risk scores the patients were divided into high and low risk **Figure 4(E)**. The KM-curve represents that patient with high risk had a poor prognosis compared to low-risk patients (Pvalue < 0.05) **Figure 4(D)**. Further, for the prediction of 3 years survivability using the ROC curve was generated with their AUC value **Figure 4(F)**. For each gene hazard ratio (HRs) and 95% confidence intervals (Cis) were analysed in forest plot **Figure 4(G)**.

**Figure 4:**
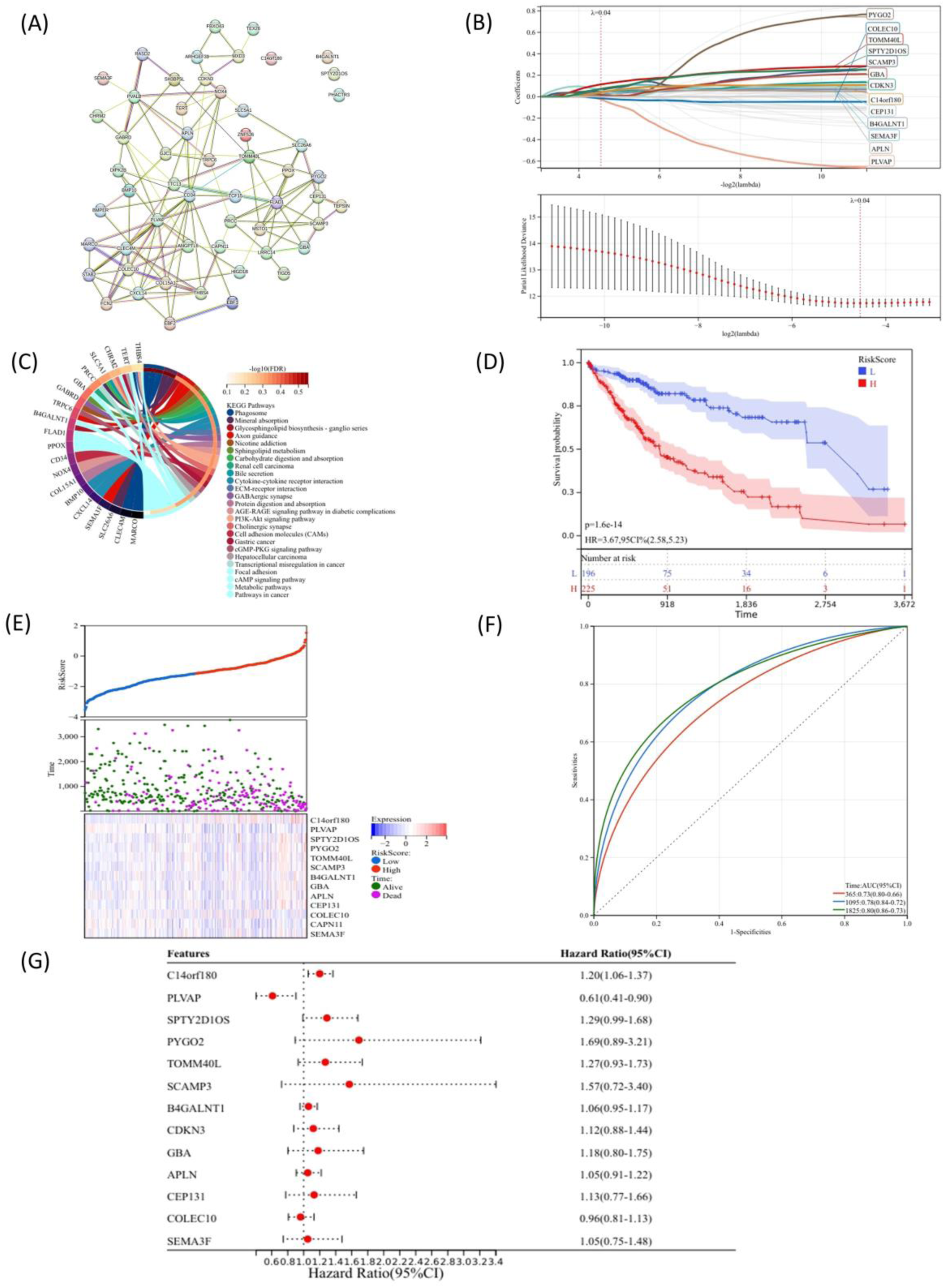
PPI interaction, KEGG pathways and prognostic value of 55 protein coding genes. (A) Protein-protein interaction of 55 protein coding genes. (B) Lasso-penalized cox regression (λ=0.04). (C) The KEGG pathways. (D)The KM-curve for high and low risk score impact on survival analysis. (E) The 13 genes with risk scores stratified into high and low risk sangerBox (http://vip.sangerbox.com/). (F) ROC curve (G) Hazard ratios with 95% CI.

### B4GALNT1 expression in other TCGA tissue

In this study, we investigated that B4GALNT1 is differentially upregulated in LIHC with pvalue < 0.05 and Kaplan-Meier Log rank P=0.002. Further, to investigate the role of B4GALNT1 we started our analysis using data from the TIMER2 website. We examined the levels of B4GALNT1 expression in various cancer types. B4GALNT1 expression was considerably higher in 19 tumour types compared to matching TCGA normal tissues, according to data collected from the TCGA database **Figure 5(A-C)**. As depicted in **Figure 5(A)**, B4GALNT1 expression was noticeably higher in tumour tissues breast invasive carcinomas (BRCA), cholangiocarcinoma (CHOL), Head and Neck squamous cell carcinoma (HNSC), Colon adenocarcinoma (COAD), Kidney renal clear cell carcinomas (KIRC), Liver hepatocellular carcinoma (LIHC), lung squamous cell carcinomas (LUSC), Lung adenocarcinoma (LUAD), Prostate adenocarcinoma (PRAD) than in the corresponding normal tissues. The same observable difference was observed in **Figure 5(B-C)** obtained from ULACAN and TNMplot. The expression of B4GALNT1 was significantly increased in LIHC tissues compared with normal samples **Figure 5(D)** and has the same result as in **Figure 5(A-C)**, with a pvalue < 0.05. The survival plot obtained from KMplot showed higher expression has poor survival **Figure 5(E)**. This explains that the expression of B4GALNT1 in LIHC is associated with survival. The expression of B4GALNT1 was compared with stage1,2,3 and 4 and it was observed significant difference of stage1,2 and 3 compared to normal samples but nor with stage 4 **Figure 5(F)**. The promoter methylation level of B4GALNT1 of cancer compared to normal sampled and a significant difference was observed **Figure 5(G)** which suggests the role of B4GALNT1 in methylation. Additionally, we examined the relationship between B4GALNT1 expression and different clinical aspects using UALCAN, of patient’s race, age, weight, gender, histological subtypes, tumour grades, nodal status and TP53 mutation in **Supplementary Figure S1(A-H)**. The significant difference with pvalue marked in red colour. Furthermore, the prognostic value of B4GALNT1 in different TCGA cancer was compared using the R packages such as “survminer” and “survival” with p < 0.05 for overall survival, progression free survival, disease free survival and disease specific survival in **Supplementary Figure S2(A-D)**. The expression of B4GALNT1 was associated with OS, DFS, PFS and DSS in some cancers. The significant difference in survival analysis in TCGA cancers with significant association of B4GALNT1 in survival is shown in **Supplementary Figure S3(A-D)**.

**Figure 5:**
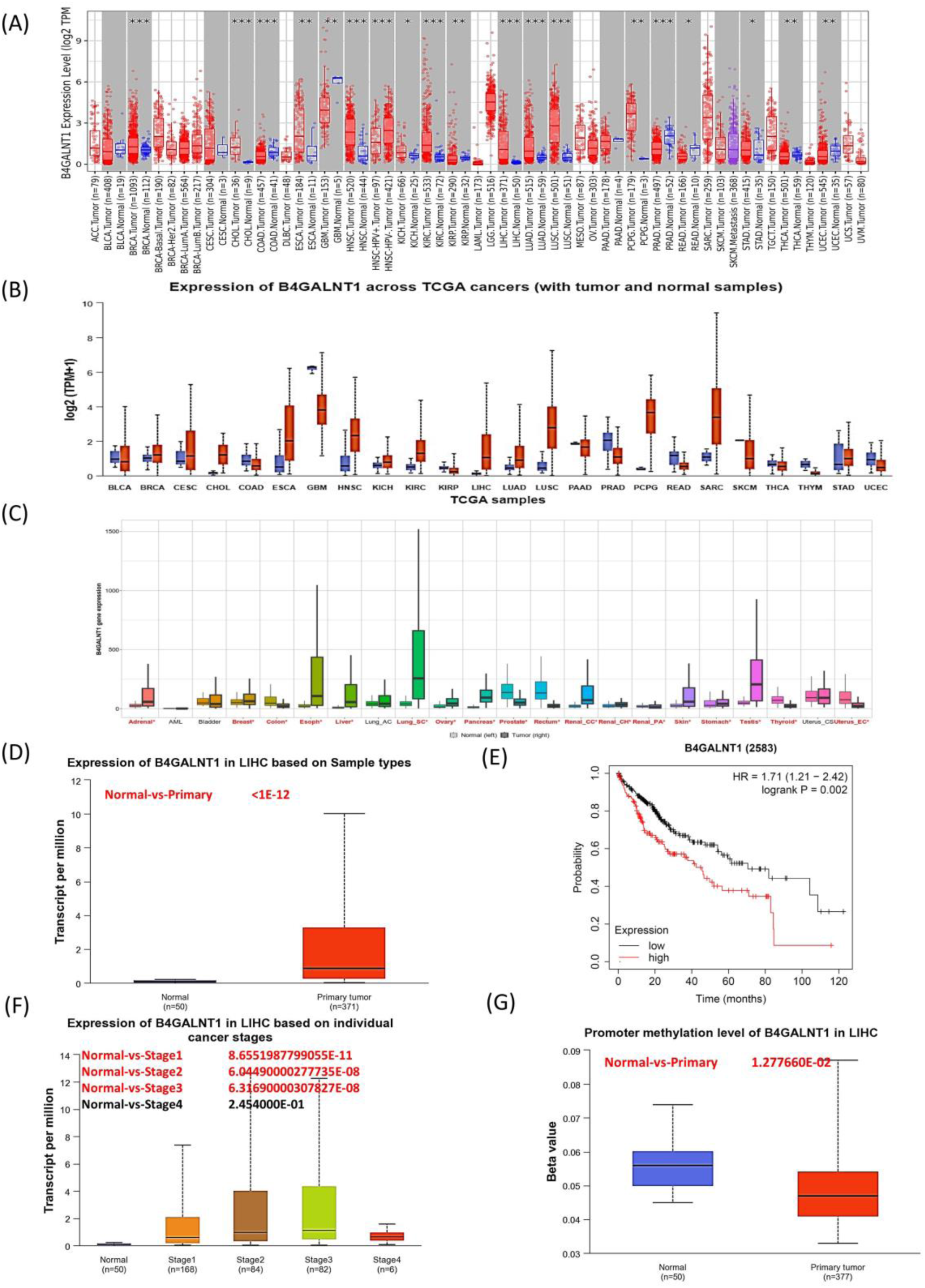
The detailed level of expression of B4GALNT1 different TCGA cancers. (A) Using TIMER2, (B) ULACAN, (C) TNM plotter, (D) Expression in LIHC, (E) Survival plot of LIHC samples, (F) Expression of B4GALNT1 in different stages, and (G) Methylation pattern of B4GALNT1.

### Expression of B4GALNT1 in other datasets, survival analysis of B4GALNT1 and neighbour genes in LIHC

The expression of B4GALNT1 were examined through CancerLivER database for affymetrix, illumina, agilent (set1 and set2) and cross checked with TCGA samples. The expression of B4GALNT1 was found to be upregulated in all the methods. **Figure 6(A)** represents the number of samples used for the analysis in tumor and adjacent tissue normal and the expression of B4GALNT1 in different NGS methods. Furthermore, B4GALNT1 survival analysis was determined using single genes-based method in KMplot for OS, RFS, PFS and DSS and showed poor survival when the expression is high **Figure 6(B-E).** To investigate the functional network of B4GALNT1 neighbouring genes in LIHC, we first used LinkedOmics to identify B4GALNT1 neighbouring genes. **Figure 6(F)** illustrates the results as a volcano plot. In addition, the heat map showed the top 50 positively and negatively associated genes. The most 5 muted genes AXIN1, DNAH10, SYNE2, TP53 and LRRI1Q associated with B4GALNT1 (p < 0.05) are shown in boxplot of mutated and wild type **Figure 6(G)**. **Figure 6(H)** clearly revealed the AUC values of B4GALNT1 in various TCGA tumours. Surprisingly, we discovered that B4GALNT1 showed high predictive value for several types of cancer. CHOL (AUC = 0.95), ESCA (AUC = 0.80), GBM (AUC = 0.87), HNSC (AUC = 0.85), KIRC (AUC = 0.86), LIHC (AUC = 0.90), LUSC (AUC = 0.93), MESO (AUC = 0.92), PCPG (AUC = 0.99), and SARC (AUC = 0.87) are among the cancer types with AUC values greater than 0.8. The findings suggested that B4GALNT1 is a good predictor of tumour prognosis. **Figure 6(I)** depicts the ROC curve for LIHC with AUC= 0.90.

**Figure 6:**
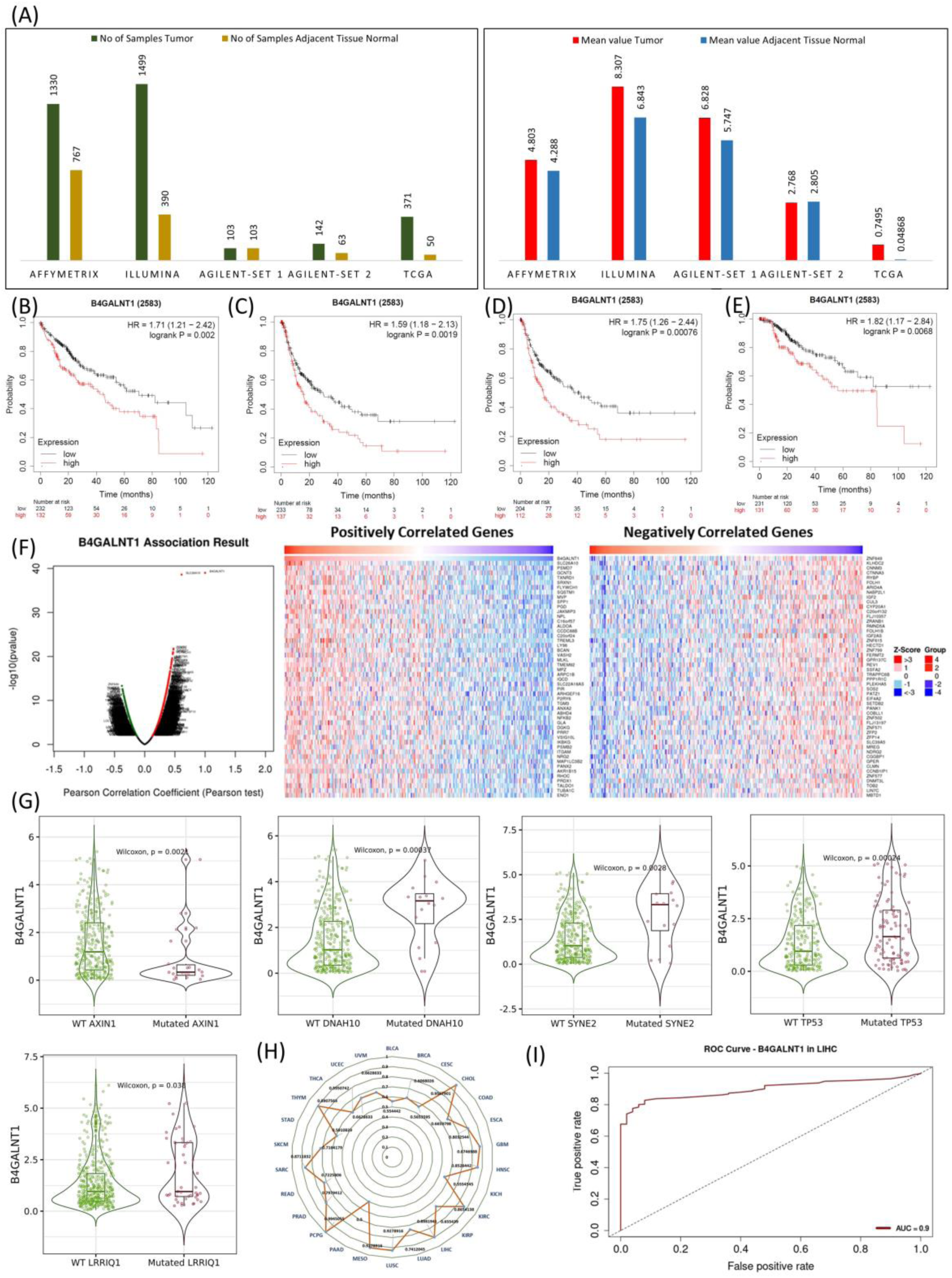
Expression of B4GALNT1 in different datasets using CancerLivER. (A) Different datasets for HCC and the mean expression of B4GALNT1 in tumor and adjacent tissue normal. (B-E) survival probability of B4GALNT1 samples in OS, RFS, PFS and DSS. (F) Volcano plot showing correlated genes with B4GALNT1, positively and negatively 50 correlated genes. (G) Association of B4GALNT1 genes with wt AXIN1, DNAH10, SYNE2, TP53, LRRIQ1 genes and mutated genes with (p < 0.05). (H) The pan-cancer AUC values for B4GALNT1 across the curve. (I) ROC plot for B4GALNT1 in LIHC with AUC = 0.9.

### Analysis of high and low B4GALNT groups in Hepatocellular Carcinoma assessing TME Score, AFP, and exploring correlations with immune cells

We looked at the relationship between B4GALNT1 and alpha fetoprotein (AFP) in LIHC patients and discovered that AFP with high expression in LIHC patients had higher expression **(Figure 7(A))**. The findings demonstrated that elevated B4GALNT1 expression in cancer was closely related to AFP levels in LIHC. We next used sangerbox (http://sangerbox.com/) to investigate the role of B4GALNT1 and immunological genes in pan-cancer **Figure 7(B)**. The results indicated a robust interaction of B4GALNT1 with both stimulatory and inhibitory immune genes across various TCGA cancers. In a surprising revelation, it was observed that CD276, TGFB1, VEGFA, HAVCR2, PDCD1, and CTLA demonstrated strong correlations with a majority of TCGA malignancies, notably LIHC, with statistical significance (p < 0.05). Regarding stimulatory immune genes, CD70, CD27, CD28, IL1B, and CD40 exhibit significant associations with the majority of TCGA malignancies (p < 0.05). Additionally, in TCGA-LIHC, B4GALNT1 expression demonstrated a positive correlation with ImmuneScore (r = 0.17, p < 0.01), ESTIMATEScore (r = 0.14, p < 0.01), and StromalScore (r = 0.08, p = 0.13). Because the ImmuneScore and ESTIMATEScore were significantly positively associated with the exception of the StromalScore, we believed that B4GALNT1 plays an important role in the tumour immune microenvironment in liver cancer **Figure 7(C-E)**. The high-B4GALNT1 group exhibited elevated TME scores in TCGA-LIHC ImmuneScore (p < 0.05), illustrated in **Figure 7(F)**. In LIHC, we assessed the variance in immune cell infiltration proportions between the high and low B4GALNT1 groups, revealing that certain immune cell fractions were higher in the high B4GALNT1 group (**Figure 7(G)**). In LIHC **Figure 7(H)**, the correlation of B4GALNT1 with immune cells and showed that it was positively correlated with B cells memory, B cells naive activated, Dendritic cells activated, T cells CD4 memory activated, and negatively correlated with NK cells activated. In LIHC, B4GALNT1 shows the strongest correlation to B cell memory. According to this study, B4GALNT1 is associated with immune cells and plays an important role in regulating the immune microenvironment in LIHC. Subsequently, we analysed the TIMER database to see if B4GALNT1 expression levels in LIHC were correlated with immune infiltration. B4GALNT1 was shown to be positively correlated with all immune cells (B cells, macrophages, neutrophils, dendritic cells, CD8+ and CD4+ T cells) with p< 0.05) **Supplementary Figure S4(A)**. In LIHC **Supplementary Figure S4(B)**, no immune cells showed any significant survival analysis accept when cumulative survival analysis was done which in turns demonstrated that immunological infiltrates were strongly associated (p < 0.05) with B4GALNT1. At last, GISTIC 2.0 was used to characterise somatic copy number variations such as deep deletion (-2), arm-level deletion (-1), diploid/normal (0), arm-level gain (1), and strong amplification (2). In LIHC **Supplementary Figure S4(C)**, box plots depict the distribution of each immune subgroup at each copy number status of B4GALNT1 where macrophages showed significant result for high amplification. The different algorithms such as TIMER, EPIC, MCPCOUNTER, XELL, TIDE, CIBERSORT, and CIBERSORT-ABS were used to investigate correlations between the infiltration level of various immune cells and the expression level of the B4GALNT1 gene in different TCGA tumours. LIHC was shown to be favourably correlated with cancer-associated fibroblasts and other immune cells **Supplementary Figure S4(D)**.

**Figure 7:**
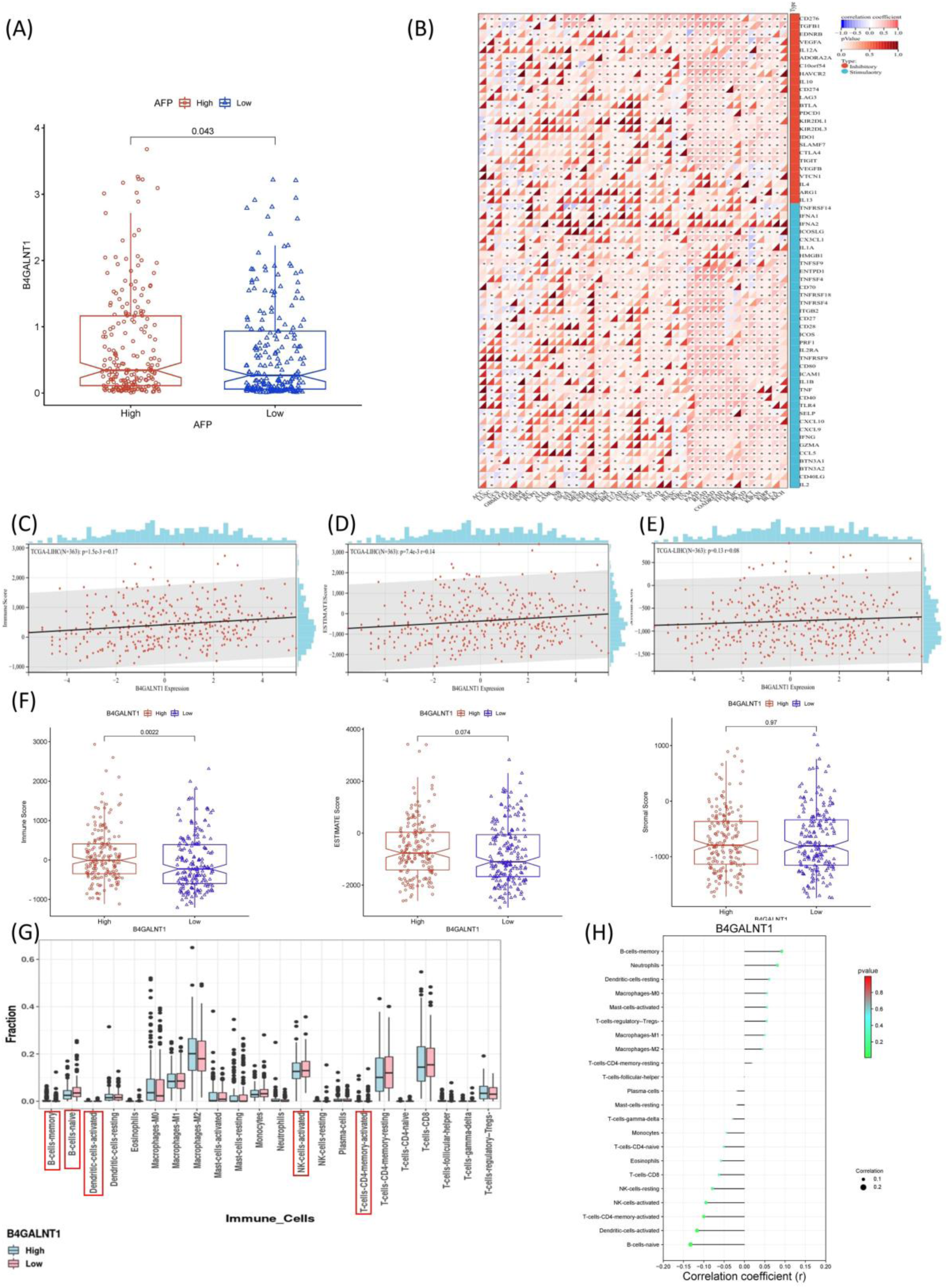
Examining the correlation between B4GALNT1 and the tumor immune microenvironment, as well as immune cells in HCC. (A) Investigating the relationship between alpha fetoprotein (AFP-High and Low) and B4GALNT1 expression. (B) Analyzing the correlation between B4GALNT1 and HLA genes in HCC, revealing a robust association. (C) Examining the correlation of B4GALNT1 with ESTIMATEScore. (D) Assessing the correlation of B4GALNT1 with ImmuneScore. (E) Investigating the correlation of B4GALNT1 with StromalScore. (F-H) Presenting boxplots of TME scores for ESTIMATEScore, ImmuneScore, and StromalScore between high−B4GALNT1 and low−B4GALNT1 groups. (I) Illustrating the difference in the fraction of immune cells between high−B4GALNT1 and low−B4GALNT1 groups, with a higher immune fraction observed in the high−B4GALNT1 group. (J) Exploring the correlation of B4GALNT1 with various cell types (p < 0.05).

### Methylation analysis of B4GALNT1

We employed the SMART (Shiny Methylation Analysis Resource Tool) App http://www.bioinfo-zs.com/smartapp for DNA methylation study, which is a user-friendly and simple-to-use online web server for extensively analysing DNA methylation data for the B4GALNT1 in LIHC investigation such as CpG visualisation, differential methylation analysis, correlation analysis, and survival analysis. The methylation pattern was observed for B4GALNT1 in LIHC for the exon, gene body and the promoter region for cancer and normal samples **Supplementary** Figure 5(A). The CpG probes were further visualized in the promoter region **Supplementary** Figure 5 **(B)**. The differentially methylation pattern among different TCGA cancer for B4GALNT1 revealed that there is a significant difference in cancer and normal samples in 15 TCGA cancers including BRCA, CHOL, COAD, ESCA, HNSC, KIRC, LIHC, LUAD, LUSC, PAAD, PRAD, READ, SKCM, THCA and UCEC (p < 0.05) **Supplementary** Figure 5 **(C).** Among CpG probes only 5 CpG probes (cg01723148, cg25392692, cg12230728, cg05771369 and cg21361094) showed significant differences when compared LIHC to normal samples. **Supplementary** Figure 5(C) indicating these CpG probes might play important roles in methylation. These probes very further analysed for stage wise difference, correlation, and survival analysis in **Supplementary** Figure 6**(A-C).**

### B4GALNT1-related gene enrichment analysis

We used the STRING tool to study 50 binding proteins and B4GALNT1 expression-related genes to know more about how the B4GALNT1 gene influences different biological activities. Text mining, experiments, databases, co-expression, Gene Fusion, and co-occurrence were used to identify these 50 proteins. The protein-protein interaction network of interacting proteins with B4GALNT1 is shown in **Figure 8(A-B)** using STRING and GeneMANIA. Further, using these 50 interacting genes KEGG, biological process, cellular component and molecular function were studied. KEGG data showed that the pathways that were significantly involved in Glycosphingolipid biosynthesis, Glycosaminoglycan biosynthesis, metabolic process other glycan related pathways. Further, GO enrichment analysis data showed that most of these genes are involved in O-glycan processing, Glycosylation, Protein glycosylation and other process. These GO and KEGG pathways were visualised using bubble plot **Figure 8(C-F)**.

**Figure 8:**
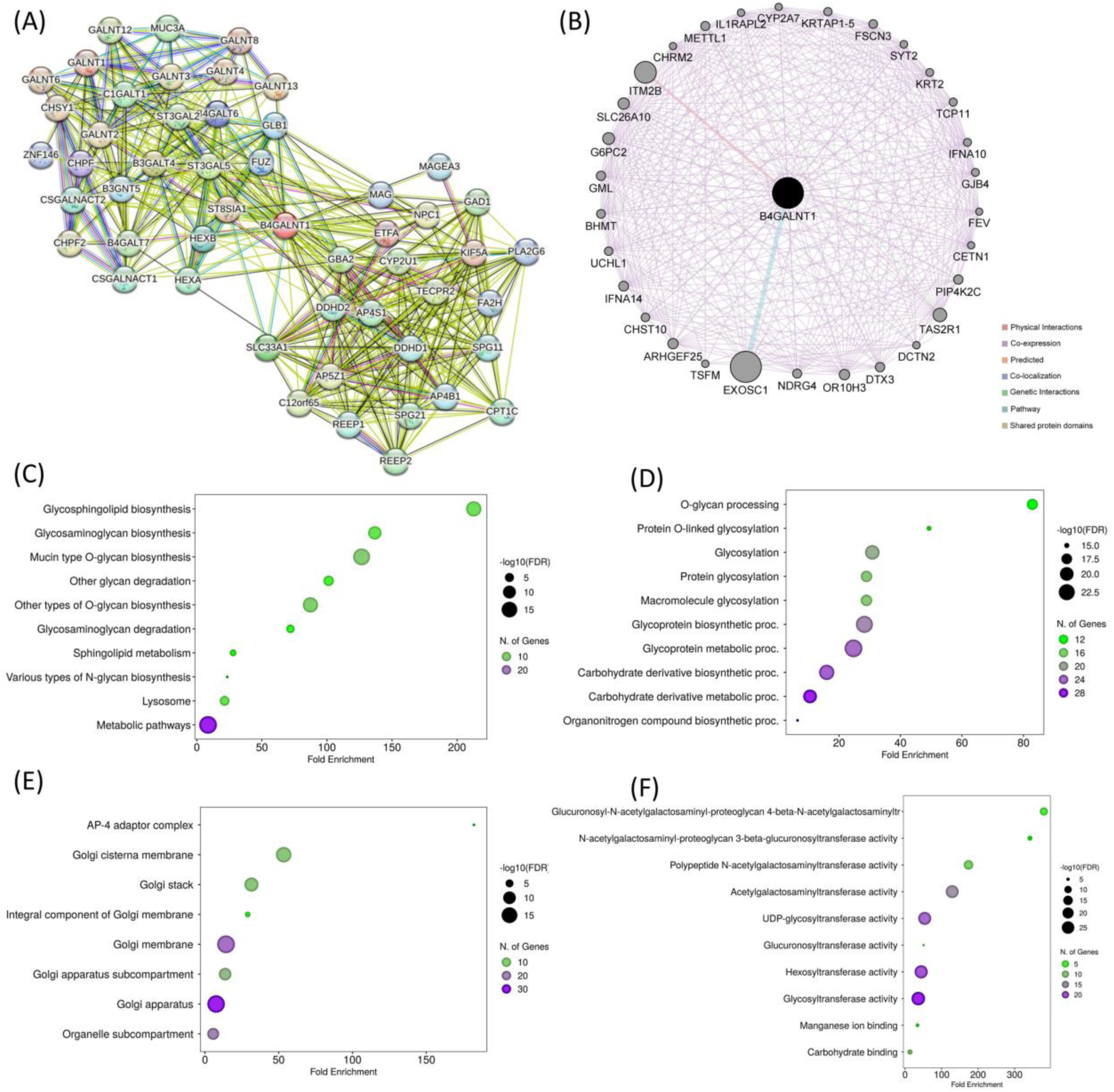
B4GALNT1related genes enrichment analysis (A) top 50 genes associted with B4GALNT1 PPI network, (B) Identifying genes that interact with B4GALNT1 in tumors through the Protein-Protein Interaction (PPI) network analysis using GeneMANIA. and (C-F) KEGG, BP, CC, MF of top 50 B4GALNT1 associated proteins.

### Gene alteration analysis

The cBioPortal dataset examined changes in B4GALNT1 mRNA expression. As indicated in **Figure 9(A)**, the most common form of change in LIHC was high of B4GALNT1, with the maximum mutational frequency reaching 2.4%. sarcoma was seen to be highest alteration frequency of B4GALNT1. The mutations landscape in different cancers for B4GALNT1 was shown in **Figure 9(B)** along with the frequency of mutation observed for liver cancers in **Figure 9(C)**. The alpha fold structure available for B4GALNT1 was used for the structural analysis with mutations in liver cancers. **Figure 9(D)** demonstrated the 3D structure of B4GALNT1 from alpha-fold model. The different mutations found in B4GALNT1 loci and corresponding cases for liver cancer from cBioportal are shown in **Figure 9(E)**. Further, the missense mutations in **Figure 9(E)** which we found from cBioportal, Cosmic Database, and Human Variant Database were filtered out from 16 different mutations which is found to be affected when analysed using in-silico approach such as SIFT, I-Mutent, Polyphen and CADD-Phred methods to 6 mutations (P64Q, D478H, P507Q, R340Q, A311S and S131F). Moreover, patients with B4GALNT1 amplification had considerably poorer progression free survival than those with no change **Supplementary Figure S7 (A-D).**

**Figure 9:**
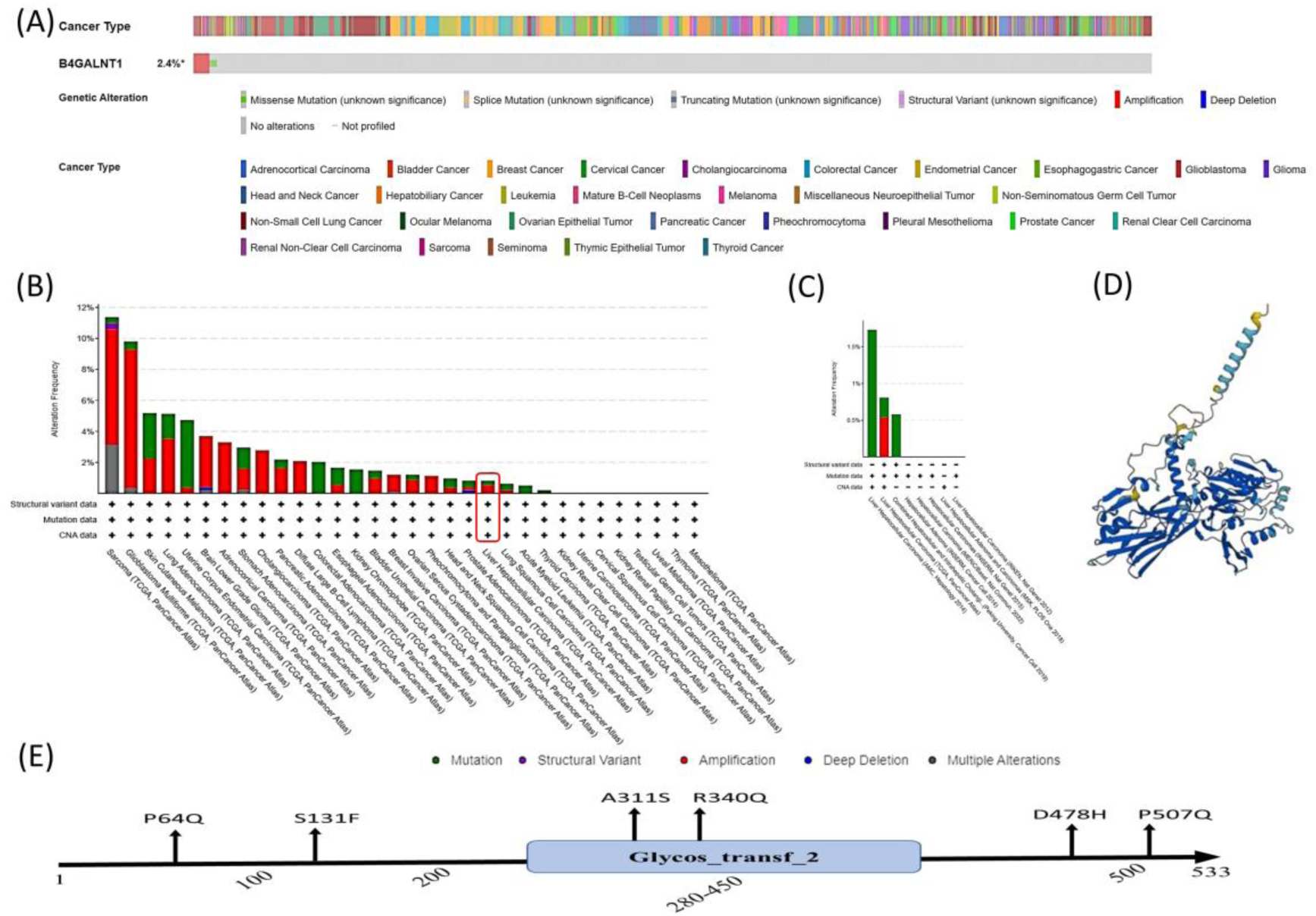
Mutational landscape of B4GALNT1 in different tumors (A) The frequency of B4GALNT1 genetic alterations in various tumour samples. (B) Variations in the copy number of B4GALNT1 across diverse tumors. Green denotes mutations, purple signifies structural variants, red indicates amplifications, blue represents deep deletions, and grey reflects multiple alterations. (C) Copy number variation of B4GALNT1 in Liver hepatocellular carcinoma (D) Predicted 3D structure of Alpha fold of B4GALNT1 (E) Mutations of B4GALNT1 in liver cancer profile; most of them are missense mutations.

### B4GALNT1 structural analysis upon mutation

**RMSD:** Through extensive all-atom molecular dynamics simulations conducted on the B4GALNT1 wild type and several of its genetic variants, the RMSDs were computed. These RMSD values were determined by aligning the molecular trajectories with their respective initial conformations, which served as the input for the MD simulations. Based on the RMSD plots for both C-α atoms (**Figure 10 (A)**) and side-chain atoms (**Figure 10 (C)**) of B4GALNT1, it is evident that the various B4GALNT1 variants exhibit greater conformational stability when compared to the wild-type B4GALNT1 structure. These plots indicate that the structural deviations in the variants are significantly minimized, highlighting the enhanced structural stability inherent in these genetic variants in contrast to the wild-type B4GALNT1 structure. **Supplementary Table S4** provides a summary of the average RMSDs for the B4GALNT1 wild type and its various variants. The RMSD data demonstrates that mutations at specific amino acid positions within B4GALNT1 have a significant impact on the protein’s structural stability, surpassing that of the wild type. This observation suggests that these genetic mutations lead to a more stable conformation in the B4GALNT1 protein, underlining the structural consequences of these specific amino acid alterations.

**Figure 10:**
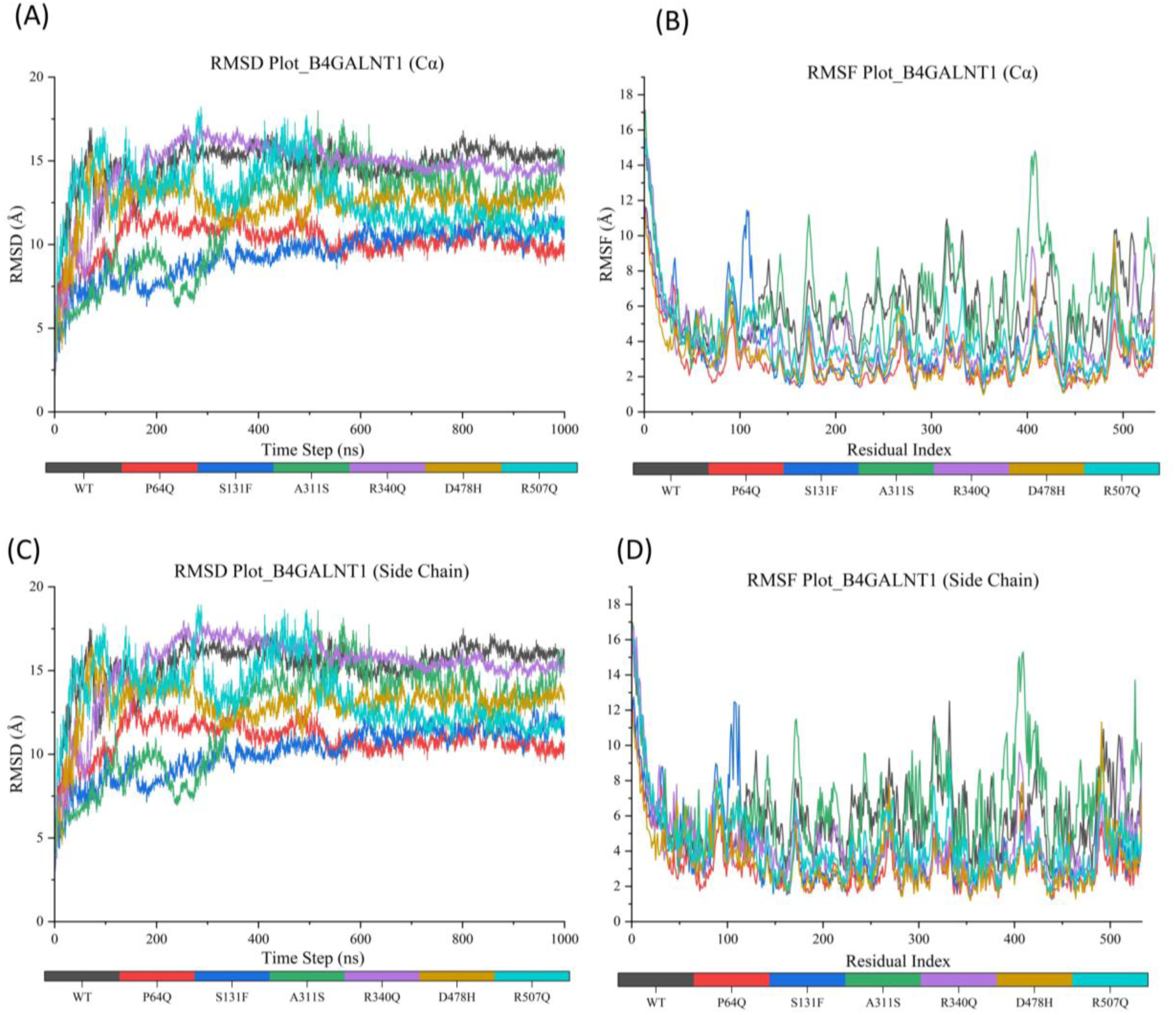
MD simulation study of B4GALNT1 WT and different mutation sites (A) Root-Mean-Square Deviation (RMSD) Analysis of B4GALNT1 and Its Variants (C-α atoms) during the 1μs of MD simulation. (B) Root-Mean-Square Fluctuation (RMSF) Analysis of B4GALNT1 and Its Variants (C-α atoms) during the 1μs of MD simulation., (C) Root-Mean-Square Deviation (RMSD) Analysis of B4GALNT1 and Its Variants (Side Chain atoms) during the 1μs of MD simulation, and (D) Root-Mean-Square Fluctuation (RMSF) Analysis of B4GALNT1 and Its Variants (Side Chain atoms) during the 1μs of MD simulation.

The Wild Type B4GALNT1 has an RMSD of 14.705 Å with a standard deviation of approximately 1.50 Å, indicating that the structure of the protein exhibits some degree of deviation during the simulation. When comparing this to the B4GALNT1 variants, several noteworthy observations can be made. The P64Q variant shows the lowest RMSD at 10.241 Å, suggesting that this mutation results in a more stable protein structure. Similarly, the S131F variant exhibits a low RMSD of 9.417 Å, indicating a high degree of structural stability. However, variants A311S, D478H, R340Q, and R507Q all show RMSD values higher than the Wild Type, with notable differences in RMSD for A311S (11.752 Å) and R340Q (14.53 Å).

**Figure 10 (A)** in the analysis indicates that the B4GALNT1 protein with the P64Q variant exhibits a distinctive structural behaviour. It appears to adopt two convergent equilibrium states during the course of the MD simulation. The first equilibrium phase is noticeable within the time frame of 170 nanoseconds to 500 nanoseconds. Subsequently, a second equilibration phase begins at 550 nanoseconds and persists throughout the production run, which extends to 1 microsecond. This second phase is characterized by lower structural deviation, as evidenced by the averaged C-α Root-Mean-Square Deviation (RMSD) of 10.241 ±1.01 Å. Furthermore, a noteworthy observation in the analysis is that after the initial 500 nanoseconds of the molecular dynamics (MD) simulation, all the B4GALNT1 variants, including the wild type, demonstrate a similar level of conformational stability. The structural behaviour appears to converge among the variants during this phase. Particularly, the B4GALNT1 R340Q variant exhibits a thermodynamic behaviour resembling that of the wild type B4GALNT1, with no significant changes in its structural characteristics observed. This suggests that, after an initial period, the R340Q variant returns to a conformational state comparable to the wild type, indicating that this specific mutation does not induce substantial alterations in the protein’s stability. These findings suggest that specific mutations in the B4GALNT1 gene can have a significant impact on the protein’s structural integrity, which can, in turn, have implications for its function and interactions within biological systems.

**RMSF: Supplementary Table S5** also provides RMSF data for different variants of the B4GALNT1 protein, specifically focusing on the fluctuations in the C-α atoms and side chain atoms. RMSF is a metric used to measure the variability or flexibility of individual atoms within a protein’s structure. The RMSF data reveals interesting insights into the dynamic behaviour of these B4GALNT1 variants. The wild-type B4GALNT1 exhibits a C-α RMSF of 5.72 Å ±1.91 Å, indicating a moderate level of atom fluctuation within the protein structure. Comparing this to the variants, it is clear that several of them display reduced RMSF values, signifying decreased atomic fluctuations and increased structural stability. The P64Q variant, for instance, demonstrates the lowest C-α RMSF at 2.831 Å, while the S131F variant shows a C-α RMSF of 3.535 Å. These lower RMSF values in P64Q and S131F suggest that these mutations lead to a more rigid protein structure with less atomic movement.

On the other hand, the A311S, D478H, R340Q, and R507Q variants have higher RMSF values compared to the Wild Type. Among them, the A311S variant has the highest RMSF at 6.099 Å. These higher RMSF values indicate increased atomic fluctuations and potentially a less stable structure in these variants. Comparing C-α RMSF to side chain RMSF values, it’s evident that in most cases, the C-α RMSF values are relatively consistent with the side chain RMSF values. This suggests that the fluctuations in the backbone atoms (C-α) are in line with the fluctuations in the side chain atoms, indicating that structural stability or flexibility affects the entire protein structure consistently.

The RMSF data highlights variations in the flexibility and stability of the B4GALNT1 variants. The P64Q and S131F variants exhibit reduced atomic fluctuations (**Figure 10 (B)** and **Figure 10 (D)**, suggesting enhanced structural stability. In contrast, the A311S, D478H, R340Q, and R507Q variants display increased atomic fluctuations, indicating a higher degree of structural flexibility. These findings shed light on the impact of specific mutations on the dynamic behaviour of the B4GALNT1 protein.

Supplementary Table S4. presents a comparison between the RMSF values of both the C-α atoms and the side chain atoms for the wild-type B4GALNT1 and the respective mutant residues. The RMSF comparison of single mutated residues with their corresponding wild-type residues provides valuable insights into the impact of these mutations on protein flexibility. Variants like P64Q, S131F, and R340Q exhibit reduced RMSF values, indicating enhanced structural stability. On the other hand, A311S and R507Q show increased RMSF values, suggesting greater flexibility and potential instability. D478H demonstrates lower RMSF values, pointing toward increased stability. These findings help in understanding the structural consequences of specific mutations in the B4GALNT1 protein.

## Discussion

There is a significant morbidity and death rates associated with liver hepatocellular carcinoma (LIHC), therefore improving clinical diagnosis and treatment of LIHC is important [45]. Several additional research have reported on the use of gene expression data to find LIHC biomarkers [46, 47]. The main disadvantage is the lack of high sample size in publicly available datasets. As a result, we sought to solve the issue of utilising the same set of samples for machine learning classification by up-sampling TCGA-LIHC data using SmoteTomek. As machine learning techniques emerge as a better method to unravelling these deep patterns in gene expression datasets in order to uncover possible new biomarkers [48-50]. However, a large number of protein coding genes in comparison to a smaller subset of samples is a barrier to improving ML classifier performance. To overcome this constraint, we used a variety of feature selection algorithms. The reason for using different feature selection methods is for two reasons: first, it aids in identifying the core representative and functional genes that can potentially distinguish two cancer and normal samples, and second, one feature selection method can provide a set of genes that may differ from another feature selection technique. Following that, we used the polling approach to identify the best features that were shared by all feature selection methods. The polling-based method yielded genes, and the majority of which are known to be associated with LIHC.

Here we have evaluated the performance of eleven ML-based classification algorithms viz.Random Forest (RF), K-Nearest Neighbours (KNN), Decision Tree (DT), Support Vector Machine (SVM), XGBoost (XGB), Naive Bayes (NB), AdaBoost (AB), Quadratic Discriminant Analysis (QDA), Multi-layer Perceptron (MLP), ExtraTrees (ET), and Logistic Regression (LR). These algorithms were used in this study to develop binary classification models for classification of LIHC and normal. The analysis found that ML algorithms could more accurately classify LIHC. Almost every ML algorithm improved sensitivity and specificity.

As indicated in the results section, Naive Bayes Classifiers outperform all other classifiers in terms of accuracy and AUC for the whole dataset trained with the 55 features **Figure 3(D)**. Several genes have established roles in LIHC out of the 55 PC genes given in the results section, validated using the above-mentioned pipeline. DGE (Differential gene expression) analysis was performed on the whole dataset, and 36 PC genes were identified to be differentially expressed from ML identified genes **(Figure 3(F))**. After that using lasso penalized coz regression analysis theses 55 PC genes were reduced to 13 genes SEMA3F [51], B4GALNT1 [7], COLEC10 [52], CEP131 [53], APLN [54], GBA [55], SCAMP3 [56], TOMM40L [57], PYGO2 [58] and PLVAP [59] has been reported in literature while CAPN11, SPTY2D1OS and C14orf180 is not so far has been studied in LIHC. Additionally, these 13 genes from survival analysis predicted to be prognostic genes for OS of LIHC cases. Among which COLEC10, C14orf180 were downregulated in our study and other genes were upregulated.

SEMA3F belongs to the semaphorin protein family, which is known for its activities in cell migration, axon guidance, and angiogenesis. SEMA3F, in particular, has been investigated for its possible role in cancer. It is classified as a tumour suppressor gene in several malignancies. It promotes hepatocellular carcinoma metastasis in the liver by activating the focal adhesion pathway [60]. COLEC10 also known as Collectin Subfamily Member 10 which encodes protein coding gene and the decreased pression level of this gene has been studied as poor OS in liver cancer patients [61]. CEP131 also known as centrosomal protein 131 has oncogenic activity in different cancers including liver [62]. Apelin which is APLN found to be upregulated promoted cancer in liver through PI3/Akt signalling pathway [63]. The role of Glucosylceramidase beta (GBA) has not been studied well in liver cancer and in our study the expression of GBA found to be upregulated. SCAMP3, a membrane-trafficking protein of the secretory carrier membrane proteins (SCAMPs) family, is involved in endosome transport. And the higher indicates a poor progression in hepatocellular carcinoma and our study also found that SCAMP3 is upregulated in LIHC [64]. TOMM40L (Translocase of Outer Mitochondrial Membrane 40 Like) is a mitochondrion related gene. The role of this gene in LIHC has not been studied well but the function of this gene is reported as cellular homeostasis, apoptosis, lipid synthesis, and energy metabolism [65]. PYGO2 also known as Pygopus Family PHD Finger 2 and the over expression of PYGO2 found in malignancy such as ovarian, lung and breast [66]. In liver by decreasing E-cadherin expression it promotes metastasis and invasion in liver cancer. Plasmalemmal Vesicle Associated Protein (PLVAP) and found as potential therapeutic target for treatment of HCC [67].

The KEGG pathways analysis of gene were associated with phagosome, mineral absorption, glycosphingolipid biosynthesis, bile secretion, PI3-Akt signalling, hepatocellular carcinoma and pathways in cancer implying these differential genes have role in LIHC but we didn’t find any GO **(Figure 5(C))**.

Furthermore, we found B4GALNT1 as one of the 13 genes is an enzyme that is encoded by the gene B4GALNT1. It plays an important role in the development and normal function of the central nervous system [68]. The pathway of glycosphingolipid biosynthesis and Sphingolipid metabolism are two of its related pathways in B4GALNT1 and related protein genes (**Figure 9(C)**).

Moreover, the role of B4GALNT1 expression in the development and prognosis on LIHC has been studied recently [7]. Therefore, here we performed comprehensive and structural analysis of B4GALNT1. In our study, we found the expression of B4GALNT1 is considerably higher in LIHC after examining the Timer, UALCAN, and HCCDB databases. Further, in subgroup analyses based on stages, tumour grade, age, gender, race, and weight, the transcriptional level of B4GALNT1 was considerably higher in LIHC patients than in normal. It was observed that the frequencies of alteration in LIHC and survival analyses for B4GALNT1 were investigated using the cBioPortal.

Through long-scale molecular dynamics simulations, we have investigated the dynamic behaviour of B4GALNT1 and its mutants. The RMSD analysis reveals that genetic variants generally exhibit increased conformational stability compared to the wild type, with mutations like P64Q and S131F displaying notably lower RMSD values. The unique structural behaviour of the P64Q mutant, adopting two dynamic equilibrium states, and the convergence of stability among mutants after an initial period. The RMSF data further underscores the impact of mutations on flexibility, with P64Q and S131F showing reduced atomic fluctuations, indicative of enhanced structural stability, while A311S, D478H, R340Q, and R507Q exhibit increased fluctuations, suggesting greater flexibility. MD simulation results collectively contribute to a comprehensive understanding of how specific mutations influence the dynamic behaviour of B4GALNT1 protein, shedding light on potential functional implications within biological systems.

Our findings proved the feasibility of employing a polling-based feature selection technique with machine learning to identify biomarkers that discriminate cancer samples from normal ones. Although more data on a wider scale is required to confirm our findings, we have painstakingly assessed and analysed our results at every step, in accordance with previous related research.

## Conclusion

Our study reveals a panel of novel genes that might be considerable for therapeutic for LIHC. These set of genes found to play important role in different pathways associated with cancer. In future, these panel can be used for target sequencing which might reduce the cost of sequencing and identifying the cancer type which in turn can help to study cancer progression and personalised treatment. Our findings suggest that B4GALNT1 were involved in tumor progression which instead suggest that it could be potential therapeutic target in LIHC. Furthermore, extensive research and clinical studies are required to validate the role of B4GALNT1 in LIHC and other malignancies.

## Availability of Data and Materials

In conducting this study, we have used mainly datasets from the publically available TCGA projects https://www.cancer.gov/about-nci/organization/ccg/research/structural-genomics/tcga and all the available online bioinformatics tools outlined in the materials and methods.

## Abbreviations

aa: Amino acid
ACC: Adrenocortical carcinoma
BLCA: Bladder Urothelial Carcinoma
BRCA: Breast invasive carcinoma
CESC: Cervical squamous cell carcinoma and endocervical adenocarcinoma
DLBC: Lymphoid Neoplasm Diffuse Large B-cell Lymphoma
GBM: Glioblastoma multiforme
HNSC: Head and Neck squamous cell carcinoma
KICH: Kidney Chromophobe
KIRC: Kidney renal clear cell carcinoma
KIRP: Kidney renal papillary cell carcinoma
LAML: Acute Myeloid Leukemia
LGG: Brain Lower Grade Glioma
LUAD: Lung adenocarcinoma
LUSC: Lung squamous cell carcinoma
OV: Ovarian serous cystadenocarcinoma
PCPG: Pheochromocytoma and Paraganglioma
PRAD: Prostate adenocarcinoma
SARC: Sarcoma
SKCM: Skin Cutaneous Melanoma
TGCT: Testicular Germ Cell Tumors
THCA: Thyroid carcinoma
THYM: Thymoma
UCEC: Uterine Corpus Endometrial Carcinoma
UCS: Uterine Carcinosarcoma
CHOL: Cholangiocarcinoma
COAD: Colon adenocarcinoma
ESCA: Esophageal carcinoma
LIHC: Liver hepatocellular carcinoma
PAAD: Pancreatic adenocarcinoma
READ: Rectum adenocarcinoma.
STAD: Stomach adenocarcinoma
INHBA: Inhibin subunit beta A
GI: Gastrointestinal
CPTAC: Clinical proteomic tumor analysis consortium
BP: Biological process
CC: Cellular component
DFS: Disease-free survival
GEPIA: Gene expression profiling interactive analysis
GO: Gene ontology
KEGG: Kyoto encyclopedia of genes and genomes
GTEx: Genotype-tissue expression
TIMER: Tumor immune estimation resource

## Author Contributions

AS, PKS and RV have designed as well as conceptualized the study. RV performed the analysis and wrote the manuscript. KBL performed the MD simulation analysis and wrote the manuscript. AS, and PKS approved further analysis, edited and reviewed the manuscript. All authors approved the manuscript. All authors contributed to the research article and approved for the submitted version.

## Acknowledgments

The researchers extend their gratitude to the Shiv Nadar Institution of Eminence (SNIoE) for furnishing the essential research framework and infrastructure vital to the successful execution of this study. The authors would like to express their gratitude to the SNIoE for providing access to the high-performance computing resource (HPC-magus02). Dr. Kiran Bharat Lokhande acknowledges SNIoE for the Post-Doctoral Fellowship.

## References

1. Sung, H., Ferlay, J., Siegel, R. L., Laversanne, M., Soerjomataram, I., Jemal, A., & Bray, F. (2021). Global cancer statistics 2020: GLOBOCAN estimates of incidence and mortality worldwide for 36 cancers in 185 countries. CA: a cancer journal for clinicians, 71(3), 209-249.

2. He, Q., Fan, B., Du, P., & Jin, Y. (2022). Construction and Validation of Two Hepatocellular Carcinoma-Progression Prognostic Scores Based on Gene Set Variation Analysis. Frontiers in Cell and Developmental Biology, 10, 806989.

3. Balogh, Julius, et al. "Hepatocellular carcinoma: a review." Journal of hepatocellular carcinoma (2016): 41–53.

4. Arbuthnot, P., & Kew, M. (2001). Hepatitis B virus and hepatocellular carcinoma. International journal of experimental pathology, 82(2), 77–100.

5. Brancato, V., Garbino, N., Salvatore, M., & Cavaliere, C. (2022). MRI-based radiomic features help identify lesions and predict histopathological grade of Hepatocellular carcinoma. Diagnostics, 12(5), 1085.

6. Weinstein, J. N., Collisson, E. A., Mills, G. B., Shaw, K. R., Ozenberger, B. A., Ellrott, K., … & Stuart, J. M. (2013). The cancer genome atlas pan-cancer analysis project. Nature genetics, 45(10), 1113–1120.

7. Liu, W., Chen, Y., Yang, J., Guo, M., & Wang, L. (2023). B4GALNT1 promotes carcinogenesis by regulating epithelial–mesenchymal transition in hepatocellular carcinoma based on pan-cancer analysis. The Journal of Gene Medicine, e3552.

8. Groux-Degroote, S., Guérardel, Y., & Delannoy, P. (2017). Gangliosides: structures, biosynthesis, analysis, and roles in cancer. Chembiochem, 18(13), 1146–1154.

9. Yoshida, H., Koodie, L., Jacobsen, K., Hanzawa, K., Miyamoto, Y., & Yamamoto, M. (2020). B4GALNT1 induces angiogenesis, anchorage independence growth and motility, and promotes tumorigenesis in melanoma by induction of ganglioside GM2/GD2. Scientific reports, 10(1), 1199.

10. Yang, H., Li, W., Lv, Y., Fan, Q., Mao, X., Long, T., … & Zhang, H. (2019). Exploring the mechanism of clear cell renal cell carcinoma metastasis and key genes based on multi-tool joint analysis. Gene, 720, 144103.

11. Che, M. I., Huang, J., Hung, J. S., Lin, Y. C., Huang, M. J., Lai, H. S., … & Huang, M. C. (2014). β1, 4-N-acetylgalactosaminyltransferase III modulates cancer stemness through EGFR signaling pathway in colon cancer cells. Oncotarget, 5(11), 3673.

12. Jiang, T., Wu, H., Lin, M., Yin, J., Tan, L., Ruan, Y., & Feng, M. (2021). B4GALNT1 promotes progression and metastasis in lung adenocarcinoma through JNK/c-Jun/Slug pathway. Carcinogenesis, 42(4), 621–630.

13. Jing, S., Deng, Z., Liang, L., & Liang, J. (2020). B4GALNT1 enhances cell proliferation and growth in oral squamous cell carcinoma via p38 and JNK MAPK pathway. Translational Cancer Research, 9(4), 2340.

14. Liang, Y. J., Ding, Y., Levery, S. B., Lobaton, M., Handa, K., & Hakomori, S. I. (2013). Differential expression profiles of glycosphingolipids in human breast cancer stem cells vs. cancer non-stem cells. Proceedings of the National Academy of Sciences, 110(13), 4968–4973.

15. Danolic, D., Heffer, M., Wagner, J., Skrlec, I., Alvir, I., Mamic, I., … & Puljiz, M. (2020). Role of ganglioside biosynthesis genetic polymorphism in cervical cancer development. Journal of Obstetrics and Gynaecology, 40(8), 1127–1132.

16. Dad, R., Malik, U., Javed, A., Minassian, B. A., & Hassan, M. J. (2017). Structural annotation of Beta-1, 4-N-acetyl galactosaminyltransferase 1 (B4GALNT1) causing Hereditary Spastic Paraplegia 26. Gene, 626, 258-263.

17. Wu, G., Lu, Z. H., Seo, J. H., Alselehdar, S. K., DeFrees, S., & Ledeen, R. W. (2020). Mice deficient in GM1 manifest both motor and non-motor symptoms of Parkinson’s disease; successful treatment with synthetic GM1 ganglioside. Experimental Neurology, 329, 113284.

18. Hong, J. M., Jeon, H., Choi, Y. C., Cho, H., Hong, Y. B., & Park, H. J. (2021). A compound heterozygous pathogenic variant in B4GALNT1 is associated with axonal Charcot-Marie-tooth disease. Journal of Clinical Neurology (Seoul, Korea), 17(4), 534.

19. Gao, J., Aksoy, B. A., Dogrusoz, U., Dresdner, G., Gross, B., Sumer, S. O., … & Schultz, N. (2013). Integrative analysis of complex cancer genomics and clinical profiles using the cBioPortal. Science signaling, 6(269), pl1-pl1.

20. Bamford, S., Dawson, E., Forbes, S., Clements, J., Pettett, R., Dogan, A., … & Wooster, R. (2004). The COSMIC (Catalogue of Somatic Mutations in Cancer) database and website. British journal of cancer, 91(2), 355–358.

21. Ganesan, K., Kulandaisamy, A., Binny Priya, S., & Gromiha, M. M. (2019). HuVarBase: A human variant database with comprehensive information at gene and protein levels. PLoS One, 14(1), e0210475.

22. Robinson, M. D., McCarthy, D. J., & Smyth, G. K. (2010). edgeR: a Bioconductor package for differential expression analysis of digital gene expression data. bioinformatics, 26(1), 139–140.

23. Goel, G., Maguire, L., Li, Y., & McLoone, S. (2013). Evaluation of sampling methods for learning from imbalanced data. In Intelligent Computing Theories: 9th International Conference, ICIC 2013, Nanning, China, July 28-31, 2013. Proceedings 9 (pp. 392-401). Springer Berlin Heidelberg.

24. Smola, A. J., & Schölkopf, B. (1998). Learning with kernels (Vol. 4). GMD-Forschungszentrum Informationstechnik.

25. Powers, D. M. W. (2011). Evaluation: From precision, recall and f-measure to roc., informedness, markedness.

26. Love, M. I., Huber, W., & Anders, S. (2014). Moderated estimation of fold change and dispersion for RNA-seq data with DESeq2. Genome biology, 15(12), 1–21.

27. Lánczky, A., & Győrffy, B. (2021). Web-based survival analysis tool tailored for medical research (KMplot): development and implementation. Journal of medical Internet research, 23(7), e27633.

28. Kaur, H., Bhalla, S., Kaur, D., & Raghava, G. P. (2020). CancerLivER: a database of liver cancer gene expression resources and biomarkers. Database, 2020, baaa012.

29. Li, T., Fu, J., Zeng, Z., Cohen, D., Li, J., Chen, Q., … & Liu, X. S. (2020). TIMER2. 0 for analysis of tumor-infiltrating immune cells. Nucleic acids research, 48(W1), W509-W514.

30. Chandrashekar, D. S., Bashel, B., Balasubramanya, S. A. H., Creighton, C. J., Ponce-Rodriguez, I., Chakravarthi, B. V., & Varambally, S. (2017). UALCAN: a portal for facilitating tumor subgroup gene expression and survival analyses. Neoplasia, 19(8), 649–658.

31. Bartha, Á., & Győrffy, B. (2021). TNMplot. com: a web tool for the comparison of gene expression in normal, tumor and metastatic tissues. International journal of molecular sciences, 22(5), 2622.

32. Vasaikar, S. V., Straub, P., Wang, J., & Zhang, B. (2018). LinkedOmics: analyzing multi-omics data within and across 32 cancer types. Nucleic acids research, 46(D1), D956–D963.

33. Kleinbaum, D. G., Klein, M., Kleinbaum, D. G., & Klein, M. (2012). Kaplan-Meier survival curves and the log-rank test. Survival analysis: a self-learning text, 55-96.

34. Menyhárt, O., Nagy, Á., & Győrffy, B. (2018). Determining consistent prognostic biomarkers of overall survival and vascular invasion in hepatocellular carcinoma. Royal Society open science, 5(12), 181006.

35. Shen, W., Song, Z., Zhong, X., Huang, M., Shen, D., Gao, P., … & Song, X. (2022). Sangerbox: A comprehensive, interaction-friendly clinical bioinformatics analysis platform. Imeta, 1(3), e36.

36. Gao, J., Aksoy, B. A., Dogrusoz, U., Dresdner, G., Gross, B., Sumer, S. O., … & Schultz, N. (2013). Integrative analysis of complex cancer genomics and clinical profiles using the cBioPortal. Science signaling, 6(269), pl1-pl1.

37. Forbes, S., Clements, J., Dawson, E., Bamford, S., Webb, T., Dogan, A., … & Stratton, M. R. (2006). COSMIC 2005. British journal of cancer, 94(2), 318–322.

38. Ganesan, K., Kulandaisamy, A., Binny Priya, S., & Gromiha, M. M. (2019). HuVarBase: A human variant database with comprehensive information at gene and protein levels. PLoS One, 14(1), e0210475.

39. Mostafavi, S., Ray, D., Warde-Farley, D., Grouios, C., & Morris, Q. (2008). GeneMANIA: a real-time multiple association network integration algorithm for predicting gene function. Genome biology, 9, 1–15.

40. Szklarczyk, D., Gable, A. L., Nastou, K. C., Lyon, D., Kirsch, R., Pyysalo, S., … & von Mering, C. (2021). The STRING database in 2021: customizable protein–protein networks, and functional characterization of user-uploaded gene/measurement sets. Nucleic acids research, 49(D1), D605–D612.

41. Ge, S. X., Jung, D., & Yao, R. (2020). ShinyGO: a graphical gene-set enrichment tool for animals and plants. Bioinformatics, 36(8), 2628–2629.

42. UniProt Consortium. (2015). UniProt: a hub for protein information. Nucleic acids research, 43(D1), D204–D212.

43. Bowers, K. J., Chow, D. E., Xu, H., Dror, R. O., Eastwood, M. P., Gregersen, B. A., Klepeis, J. L., Kolossvary, I., Moraes, M. A., Sacerdoti, F. D., Salmon, J. K., Shan, Y., & Shaw, D. E. (2006). Scalable algorithms for molecular dynamics simulations on commodity clusters, SC ’06 [Paper presentation]. Proceedings of the 2006 ACM/IEEE Conference on Supercomputing, Tampa, FL (pp. 43). 10.1145/1188455.1188544

44. Kollar, J., & Frecer, V. (2017). How accurate is the description of ligand-protein interactions by a hybrid QM/MM approach?. Journal of molecular modeling, 24(1), 11. 10.1007/s00894-017-3537-z

45. Ouyang, X., Fan, Q., Ling, G., Shi, Y., & Hu, F. (2020). Identification of diagnostic biomarkers and subtypes of liver hepatocellular carcinoma by multi-omics data analysis. Genes, 11(9), 1051.

46. Kaur, H., Bhalla, S., & Raghava, G. P. (2019). Classification of early and late stage liver hepatocellular carcinoma patients from their genomics and epigenomics profiles. PloS one, 14(9), e0221476.

47. Huo, Q., Ma, Y., Yin, Y., & Qin, G. (2021). Biomarker identification for liver hepatocellular carcinoma and cholangiocarcinoma based on gene regulatory network analysis. Current Bioinformatics, 16(1), 31–43.

48. Yuan, F., Lu, L., & Zou, Q. (2020). Analysis of gene expression profiles of lung cancer subtypes with machine learning algorithms. Biochimica et Biophysica Acta (BBA)-Molecular Basis of Disease, 1866(8), 165822.

49. Sinkala, M., Mulder, N., & Martin, D. (2020). Machine learning and network analyses reveal disease subtypes of pancreatic cancer and their molecular characteristics. Scientific reports, 10(1), 1212.

50. Tabl, A. A., Alkhateeb, A., ElMaraghy, W., Rueda, L., & Ngom, A. (2019). A machine learning approach for identifying gene biomarkers guiding the treatment of breast cancer. Frontiers in genetics, 10, 256.

51. Yang, H., Li, G., & Qiu, G. (2021). Bioinformatics analysis using ATAC-seq and RNA-seq for the identification of 15 gene signatures associated with the prediction of prognosis in hepatocellular carcinoma. Frontiers in Oncology, 11, 726551.

52. Ju, M., Jiang, L., Wei, Q., Yu, L., Chen, L., Wang, Y., … & Han, J. (2021). A immune-related signature associated with TME can serve as a potential biomarker for survival and sorafenib resistance in liver cancer. OncoTargets and therapy, 5065-5083.

53. Liu, X. H., Yang, Y. F., Fang, H. Y., Wang, X. H., Zhang, M. F., & Wu, D. C. (2017). CEP131 indicates poor prognosis and promotes cell proliferation and migration in hepatocellular carcinoma. The International Journal of Biochemistry & Cell Biology, 90, 1–8.

54. Lin, Z. H., Zhang, J., Zhuang, L. K., Xin, Y. N., & Xuan, S. Y. (2022). Establishment of a Prognostic Model for Hepatocellular Carcinoma Based on Bioinformatics and the Role of NR6A1 in the Progression of HCC. Journal of Clinical and Translational Hepatology, 10(5), 901.

55. Chen, W., Ma, Z., Yu, L., Mao, X., Ma, N., Guo, X., … & Zhang, Y. (2022). Preclinical investigation of artesunate as a therapeutic agent for hepatocellular carcinoma via impairment of glucosylceramidase-mediated autophagic degradation. Experimental & Molecular Medicine, 54(9), 1536–1548.

56. Feng, G., He, N., Xia, H. H. X., Mi, M., Wang, K., Byrne, C. D., … & Ye, F. (2022). Machine learning algorithms based on proteomic data mining accurately predicting the recurrence of hepatitis B-related hepatocellular carcinoma. Journal of Gastroenterology and Hepatology, 37(11), 2145–2153.

57. Xu, W., Zhao, D., Huang, X., Zhang, M., Yin, M., Liu, L., … & Xu, C. (2022). The prognostic value and clinical significance of mitophagy-related genes in hepatocellular carcinoma. Frontiers in Genetics, 13, 917584.

58. Yuan, W., Zhao, H., Zhou, A., & Wang, S. (2022). Interference of EFNA4 suppresses cell proliferation, invasion and angiogenesis in hepatocellular carcinoma by downregulating PYGO2. Cancer Biology & Therapy, 23(1), 1–12.

59. Sharma, A., Seow, J. J. W., Dutertre, C. A., Pai, R., Blériot, C., Mishra, A., … & DasGupta, R. (2020). Onco-fetal reprogramming of endothelial cells drives immunosuppressive macrophages in hepatocellular carcinoma. Cell, 183(2), 377–394.

60. Ye, K., Ouyang, X., Wang, Z., Yao, L., & Zhang, G. (2020). SEMA3F promotes liver hepatocellular carcinoma metastasis by activating focal adhesion pathway. DNA and cell biology, 39(3), 474–483.

61. Zhang, B., & Wu, H. (2018). Decreased expression of COLEC10 predicts poor overall survival in patients with hepatocellular carcinoma. Cancer management and research, 2369-2375.

62. Wang, J., Yang, X., Han, S., & Zhang, L. (2020). CEP131 knockdown inhibits cell proliferation by inhibiting the ERK and AKT signaling pathways in non-small cell lung cancer. Oncology Letters, 19(4), 3145–3152.

63. Chen, H., Wong, C. C., Liu, D., Go, M. Y., Wu, B., Peng, S., … & Yu, J. (2019). APLN promotes hepatocellular carcinoma through activating PI3K/Akt pathway and is a druggable target. Theranostics, 9(18), 5246.

64. Zhang, X., Sheng, J., Zhang, Y., Tian, Y., Zhu, J., Luo, N., … & Li, R. (2017). Overexpression of SCAMP3 is an indicator of poor prognosis in hepatocellular carcinoma. Oncotarget, 8(65), 109247.

65. Chen, S., Sarasua, S. M., Davis, N. J., DeLuca, J. M., Boccuto, L., Thielke, S. M., & Yu, C. E. (2022). TOMM40 genetic variants associated with healthy aging and longevity: A systematic review. BMC geriatrics, 22(1), 667.

66. Zhang, S., Li, J., Liu, P., Xu, J., Zhao, W., Xie, C., … & Wang, X. (2015). Pygopus-2 promotes invasion and metastasis of hepatic carcinoma cell by decreasing E-cadherin expression. Oncotarget, 6(13), 11074.

67. Wang, Y. H., Cheng, T. Y., Chen, T. Y., Chang, K. M., Chuang, V. P., & Kao, K. J. (2014). Plasmalemmal Vesicle Associated Protein (PLVAP) as a therapeutic target for treatment of hepatocellular carcinoma. BMC cancer, 14(1), 1–12.

68. Yi, H., Lin, Y., Li, Y., Guo, Y., Yuan, L., & Mao, Y. (2022). Pan-Cancer Analysis of B4GALNT1 as a Potential Prognostic and Immunological Biomarker. Journal of Immunology Research, 2022.

