## Supplementary Materials for "Machine Learning-Based Identification of B4GALNT1 as a Key Player in Hepatocellular Carcinoma: A Comprehensive Bioinformatics and Structural Analysis"

***Corresponding Author Details-**

**Dr. Prashant Shrivastava**

National Heart and Lung Institute,
Imperial College London, London,
United Kingdom.
,

**Dr. Ashutosh Singh**

Associate Professor,
Translational Bioinformatics and Computational Genomics Laboratory,
Department of Life Sciences, School of Natural Sciences (SONS),
Shiv Nadar Institution of Eminence (SNIoE), Delhi NCR, India.


***Supplementary Materials:***

*Supplementary Methodology:*

### Feature Selection

Finding a set of genes that specific to disease and biological activity from a big dimensional feature set is the main goal. It might be difficult to choose a smaller, more meaningful collection of attributes while maintaining the validity of the data. As certain feature selection techniques produce excellent results for some types of data while producing unpromising results for others utilizing the strength of all feature selection processes, therefore we have used various feature selection procedure to highlight the most important aspects. Using the normalized data set for protein coding genes dataset were used for further feature selection. Further, we used rocc package in R based on receiver operating characteristics (ROC) feature selection technique to classify cancer and normal samples. This method reduces the feature from approx 20000 features to 3149 features with AUC >= 0.90. Moreover, to get a smaller set of data that can better classify we next used different feature selection methods based on variable importance from the model such as select from model which is Meta-Transformer for selecting features based on importance weights with estimators xgboost, random forest, Support Vector Machine (linear) [28], Feature Importance using Extra Tree classifier which is a feature scoring technique and some other feature selection such as Boruta which is a wrapper built around the random forest classification algorithm involving a random shadow feature set. Univariate Feature Selection (Ranked Feature Set) is a feature set based on univariate statistical test with configurable strategy. Spikeslab. Spike and slab for variable selection in linear regression models. A feature set based on the fast correlation-based filter method (FCBF), Lasso Regression and Recursive Feature Elimination (RFE) to obtain the best gene feature subset. Finally, we obtained the features that were common in all the techniques. These Features sets from the noted Feature Selection Techniques are subjected to Polling. Polling Technique strives to find a common ground between these feature selection techniques. The Ideology is if a singular feature is termed as a significant feature by one feature selection technique, then other Feature Selection techniques should recognize that feature as significant as well. In other words, all Feature Selection techniques casts their votes to its selected features, where every vote has singleton value. Later these votes for each feature are summed up to elect the best set of features by setting up a threshold. Thresholds are relative, which are adjusted based on final required set of features that needs to be sent to the next phase. In this study, we set our threshold to >=5 stating that, a feature is selected if and only if that feature is selected by at-least a minimum of >=5 feature selection techniques which narrows down the selected features. This Polling is one of the crucial steps to get rid of features that might have been selected incidentally. This technique has been implemented in the further steps as well. Finally, we obtained 55 gene features from the gene expression datasets.

**Performance evaluation metrics**

Evaluation metrics are crucial to evaluate the performance of machine learning model, and it is necessary to adopt those metrics, which serve our purpose. For classification, we have used sensitivity, specificity, accuracy, MCC, ROC/AUC, recall, and precision. Out of all ROC/AUC, MCC, sensitivity and precision provide greater insight into the model performance.

To measure models' performance, we used standard parameters, commonly used to measure classification models' performance using sensitivity, specificity, accuracy, and Matthew's correlation coefficient (MCC) using the following equations.

$$(Sensitivity) = \frac{TP}{TP + TN} * 100$$

$$(\text{Specificity}) = \frac{TN}{TN + FP} * 100$$

$$(\text{Accuracy}) = \frac{(TP + TN)}{(TP + FP + TN + FN)} * 100$$

$$(\text{AccuracyMatthew's correlation coefficient (MCC)}) = \frac{(TP * TN) - (FP * FN)}{\sqrt{(TP + FP)(TP + FN)(TN + FP)(TN + FN)}} * 100$$

Here, False positive, false negative, true positive, and true negative predictions are denoted by FP, FN, TP, and TN, respectively.

*Supplementary Tables*

**Table S1.** The performance of LIHC classification developed using 55 features where sensitivity, specificity and accuracy are in percentage.

| Classifier | Sensitivity | Specificity | Accuracy | AUC | MCC | Precision | Recall | F1 | kappa |
|---|---|---|---|---|---|---|---|---|---|
| AB | 79.22 | 87.67 | 83.33 | 0.92 | 0.67 | 0.87 | 0.79 | 0.83 | 0.67 |
| DT | 64.94 | 67.12 | 66 | 0.67 | 0.32 | 0.68 | 0.65 | 0.66 | 0.32 |
| ET | 81.82 | 91.78 | 86.67 | 0.94 | 0.74 | 0.91 | 0.82 | 0.86 | 0.73 |
| GNB | 87.01 | 95.89 | 91.33 | 0.96 | 0.83 | 0.96 | 0.87 | 0.91 | 0.83 |
| KNN | 83.12 | 94.52 | 88.67 | 0.95 | 0.78 | 0.94 | 0.83 | 0.88 | 0.77 |
| LR | 87.01 | 91.78 | 89.33 | 0.94 | 0.79 | 0.92 | 0.87 | 0.89 | 0.79 |
| MLP | 88.31 | 91.78 | 90 | 0.95 | 0.8 | 0.92 | 0.88 | 0.9 | 0.8 |
| QDA | 80.52 | 82.19 | 81.33 | 0.89 | 0.63 | 0.83 | 0.81 | 0.82 | 0.63 |
| RF | 83.12 | 94.52 | 88.67 | 0.95 | 0.78 | 0.94 | 0.83 | 0.88 | 0.77 |
| SVC | 88.31 | 93.15 | 90.67 | 0.95 | 0.79 | 0.9 | 0.89 | 0.9 | 0.79 |
| XGB | 87.01 | 93.15 | 90 | 0.95 | 0.8 | 0.93 | 0.87 | 0.9 | 0.8 |

**Table S2**. The performance of LIHC classification developed using 55 features where sensitivity, specificity and accuracy are in percentage for validation dataset.

| Classifier | Sensitivity | Specificity | Accuracy | AUC | MCC | Precision | Recall | F1 | kappa |
|---|---|---|---|---|---|---|---|---|---|
| AB | 90.79 | 85.14 | 88 | 0.93 | 0.76 | 0.86 | 0.91 | 0.88 | 0.76 |
| DT | 57.89 | 87.84 | 72.67 | 0.75 | 0.48 | 0.83 | 0.58 | 0.68 | 0.46 |
| ET | 85.53 | 90.54 | 88 | 0.95 | 0.76 | 0.9 | 0.86 | 0.88 | 0.76 |
| GNB | 89.47 | 89.19 | 89.33 | 0.97 | 0.79 | 0.89 | 0.89 | 0.89 | 0.79 |
| KNN | 88.16 | 91.89 | 90 | 0.97 | 0.8 | 0.92 | 0.88 | 0.9 | 0.8 |
| LR | 92.11 | 93.24 | 92.67 | 0.98 | 0.85 | 0.93 | 0.92 | 0.93 | 0.85 |
| MLP | 89.47 | 87.84 | 88.67 | 0.95 | 0.77 | 0.88 | 0.89 | 0.89 | 0.77 |
| QDA | 76.32 | 83.78 | 80 | 0.88 | 0.6 | 0.83 | 0.76 | 0.79 | 0.6 |
| RF | 84.21 | 91.89 | 88 | 0.95 | 0.76 | 0.91 | 0.84 | 0.88 | 0.76 |
| SVC | 90.79 | 95.95 | 93.33 | 0.95 | 0.76 | 0.89 | 0.87 | 0.88 | 0.76 |
| XGB | 85.53 | 85.14 | 85.33 | 0.93 | 0.71 | 0.86 | 0.86 | 0.86 | 0.71 |

**Table S3.** 55 gene features after ML, along with their gene expression, log2FC, pvalue, padj, Kaplan-Meier Log Rank P-value, and AUC.

| Gene | log2 FC | pvalue | padj | Gene Expression | Kaplan-Meier Log rank P (LIHC) | AUC |
|---|---|---|---|---|---|---|
| SEMA3F | 1.284 | 6.70E-39 | 3.99E-37 | Upregulated | 0.0052 | 0.991 |
| MARCO | -4.844 | 3.90E-41 | 2.72E-39 | Downregulated | 0.3053 | 0.979 |
| NOX4 | 2.852 | 9.66E-55 | 1.40E-52 | Upregulated | 0.0464 | 0.992 |

| | | | | | | |
|---|---|---|---|---|---|---|
| PHACTR3 | -2.908 | 7.37E-30 | 2.31E-28 | Downregulated | 0.0004 | 0.963 |
| SLC5A1 | -1.936 | 2.57E-05 | 6.42E-05 | Downregulated | 0.0115 | 0.912 |
| RASD2 | 2.633 | 4.98E-49 | 5.38E-47 | Upregulated | 0.1042 | 0.967 |
| PVALB | -3.663 | 4.34E-30 | 1.38E-28 | Downregulated | 0.0269 | 0.969 |
| CDKN3 | 3.919 | 1.61E-95 | 1.00E-91 | Upregulated | 0.0066 | 0.983 |
| CLEC4M | -5.585 | 3.31E-26 | 7.44E-25 | Downregulated | 6.60E-06 | 0.991 |
| THBS4 | 5.454 | 2.34E-80 | 1.46E-77 | Upregulated | 0.0478 | 0.969 |
| SCAMP3 | 1.280 | 8.24E-40 | 5.24E-38 | Upregulated | 0.0156 | 0.993 |
| MSTO1 | 1.740 | 6.78E-61 | 1.32E-58 | Upregulated | 0.0108 | 0.997 |
| TCF15 | 2.328 | 1.73E-29 | 5.29E-28 | Upregulated | 0.002 | 0.968 |
| PLVAP | 2.918 | 1.74E-99 | 1.63E-95 | Upregulated | 9.10E-05 | 0.994 |
| ANGPTL6 | -3.025 | 7.60E-54 | 1.07E-51 | Downregulated | 0.0095 | 0.983 |
| HIGD1B | 2.849 | 4.92E-57 | 7.87E-55 | Upregulated | 0.267 | 0.984 |
| B4GALNT1 | 4.303 | 3.89E-51 | 4.64E-49 | Upregulated | 0.002 | 0.911 |
| STAB2 | -4.813 | 4.75E-64 | 1.03E-61 | Downregulated | 0.0513 | 0.985 |
| ARHGEF39 | 3.175 | 6.26E-86 | 7.80E-83 | Upregulated | 0.0004 | 0.995 |
| CAPN11 | 2.100 | 4.01E-35 | 1.92E-33 | Upregulated | 0.0003 | 0.972 |
| TRPC6 | 2.114 | 6.07E-36 | 3.04E-34 | Upregulated | 0.0005 | 0.960 |
| CEP131 | 1.813 | 3.07E-54 | 4.36E-52 | Upregulated | 0.0009 | 0.993 |
| PPOX | 1.163 | 5.16E-38 | 2.88E-36 | Upregulated | 0.2051 | 0.993 |
| PRCC | 0.956 | 2.98E-30 | 9.62E-29 | FALSE | 0.005 | 0.990 |
| TTC13 | 1.408 | 2.33E-49 | 2.53E-47 | Upregulated | 0.1795 | 0.987 |
| CXCL14 | -3.565 | 1.10E-19 | 1.40E-18 | Downregulated | 0.1216 | 0.968 |
| DIPK2B | 2.216 | 7.74E-76 | 3.29E-73 | Upregulated | 0.0853 | 0.995 |
| FBXO43 | 4.047 | 6.82E-74 | 2.71E-71 | Upregulated | 0.0025 | 0.984 |
| TOMM40L | 1.791 | 4.11E-67 | 1.08E-64 | Upregulated | 0.0203 | 0.996 |
| FCN2 | -4.880 | 7.98E-40 | 5.11E-38 | Downregulated | 0.0146 | 0.988 |
| FLAD1 | 1.025 | 4.45E-29 | 1.31E-27 | Upregulated | 0.0466 | 0.996 |
| LRRC14 | 1.481 | 2.46E-45 | 2.24E-43 | Upregulated | 0.2707 | 0.998 |
| BMP10 | -4.707 | 2.67E-19 | 3.25E-18 | Downregulated | 1.20E-08 | 0.981 |
| PYGO2 | 1.221 | 2.90E-47 | 2.90E-45 | Upregulated | 0.0006 | 0.995 |
| EBF1 | 1.998 | 4.03E-33 | 1.67E-31 | Upregulated | 0.0483 | 0.964 |
| TERT | 8.551 | 4.50E-68 | 1.40E-65 | Upregulated | 0.032 | 0.983 |
| BMPER | -4.506 | 3.12E-51 | 3.74E-49 | Downregulated | 0.0008 | 0.989 |
| TEPSIN | 1.042 | 1.01E-25 | 2.15E-24 | Upregulated | 0.1978 | 0.988 |
| ZNF526 | 0.728 | 3.51E-24 | 6.51E-23 | FALSE | 0.2582 | 0.994 |
| APLN | 3.900 | 1.75E-70 | 5.65E-68 | Upregulated | 0.0028 | 0.988 |
| CD34 | 2.091 | 5.11E-65 | 1.18E-62 | Upregulated | 0.0018 | 0.981 |
| SH3BP5L | 0.873 | 4.71E-30 | 1.49E-28 | FALSE | 0.0451 | 0.962 |

| | | | | | | |
|---|---|---|---|---|---|---|
| TEX26 | -3.891 | 8.09E-18 | 8.56E-17 | Downregulated | 1.40E-08 | 0.983 |
| GBA | 1.367 | 4.74E-37 | 2.54E-35 | Upregulated | 0.0219 | 0.991 |
| TIGD5 | 1.646 | 2.03E-37 | 1.10E-35 | Upregulated | 0.0552 | 0.965 |
| CHRM2 | -3.785 | 1.87E-15 | 1.55E-14 | Downregulated | 3.00E-08 | 0.970 |
| GJC1 | 2.356 | 5.39E-44 | 4.48E-42 | Upregulated | 0.3716 | 0.983 |
| COLEC10 | -3.999 | 2.16E-38 | 1.25E-36 | Downregulated | 0.0001 | 0.951 |
| C14orf180 | -2.952 | 6.91E-09 | 2.71E-08 | Downregulated | 0.0063 | 0.997 |
| GABRD | 4.475 | 3.90E-118 | 7.29E-114 | Upregulated | 0.0507 | 0.984 |
| COL15A1 | 3.799 | 6.06E-84 | 5.39E-81 | Upregulated | 0.0001 | 0.993 |
| MXD3 | 2.786 | 2.36E-81 | 1.58E-78 | Upregulated | 0.0025 | 0.983 |
| EBF2 | 5.259 | 4.58E-79 | 2.60E-76 | Upregulated | 0.0466 | 0.996 |
| SLC26A6 | 2.660 | 1.17E-80 | 7.53E-78 | Upregulated | 0.0014 | 0.963 |
| SPTY2D1OS | 1.208 | 1.71E-15 | 1.43E-14 | Upregulated | 0.0919 | 0.964 |

**Table S4:** The average and standard deviation of RMSDs and RMSFs of the B4GALNT1 wild type and mutant type.

| **Sr. No.** | **B4GALNT1 Variants** | **RMSD (C- α) [Å]** | **RMSD (Side Chain) [Å]** | **RMSF (C- α) [Å]** | **RMSF (Side Chain) [Å]** |
|---|---|---|---|---|---|
| 1 | **Wild Type** | 14.705 | 15.328 | 5.72 | 6.028 |
| | | ±1.50 | ±1.54 | ±1.91 | ±2.01 |
| 2 | **P64Q** | 10.241 | 10.902 | 2.831 | 3.184 |
| | | ±1.01 | ±1.02 | ±1.44 | ±1.56 |
| 3 | **S131F** | 9.417 | 10.114 | 3.535 | 3.907 |
| | | ±1.46 | ±1.48 | ±1.97 | ±2.11 |
| 4 | **A311S** | 11.752 | 12.29 | 6.099 | 6.455 |
| | | ±3.070 | ±3.065 | ±2.43 | ±2.51 |
| 5 | **R340Q** | 14.53 | 15.267 | 4.05 | 4.356 |
| | | ±2.08 | ±2.14 | ±1.97 | ±2.09 |
| 6 | **D478H** | 12.402 | 12.982 | 3.095 | 3.482 |
| | | ±1.36 | ±1.38 | ±1.49 | ±1.63 |
| 7 | **R507Q** | 12.859 | 13.52 | 4.075 | 4.405 |
| | | ±1.82 | ±1.90 | ±1.84 | ±1.99 |

**Table S5:** Mean RMSF values for the specific mutations as compared to wild-type B4GALNT1.

| Sr. No. | B4GALNT1 Variants | RMSF-Wild Type (C- α) [Å] | RMSF-Wild Type (Side Chain) [Å] | RMSF-Mutants (C- α) [Å] | RMSF-Mutants (Side Chain) [Å] |
|---|---|---|---|---|---|
| 1 | **P64Q** | 4.192 | 4.549 | 3.04 | 3.838 |
| 2 | **S131F** | 7.583 | 7.866 | 2.466 | 2.583 |
| 3 | **A311S** | 6.524 | 6.288 | 5.776 | 5.635 |
| 4 | **R340Q** | 4.663 | 3.798 | 2.434 | 2.664 |
| 5 | **D478H** | 5.264 | 6.005 | 2.566 | 2.868 |
| 6 | **R507Q** | 8.576 | 8.739 | 4.443 | 5.074 |

*Supplementary Figures*

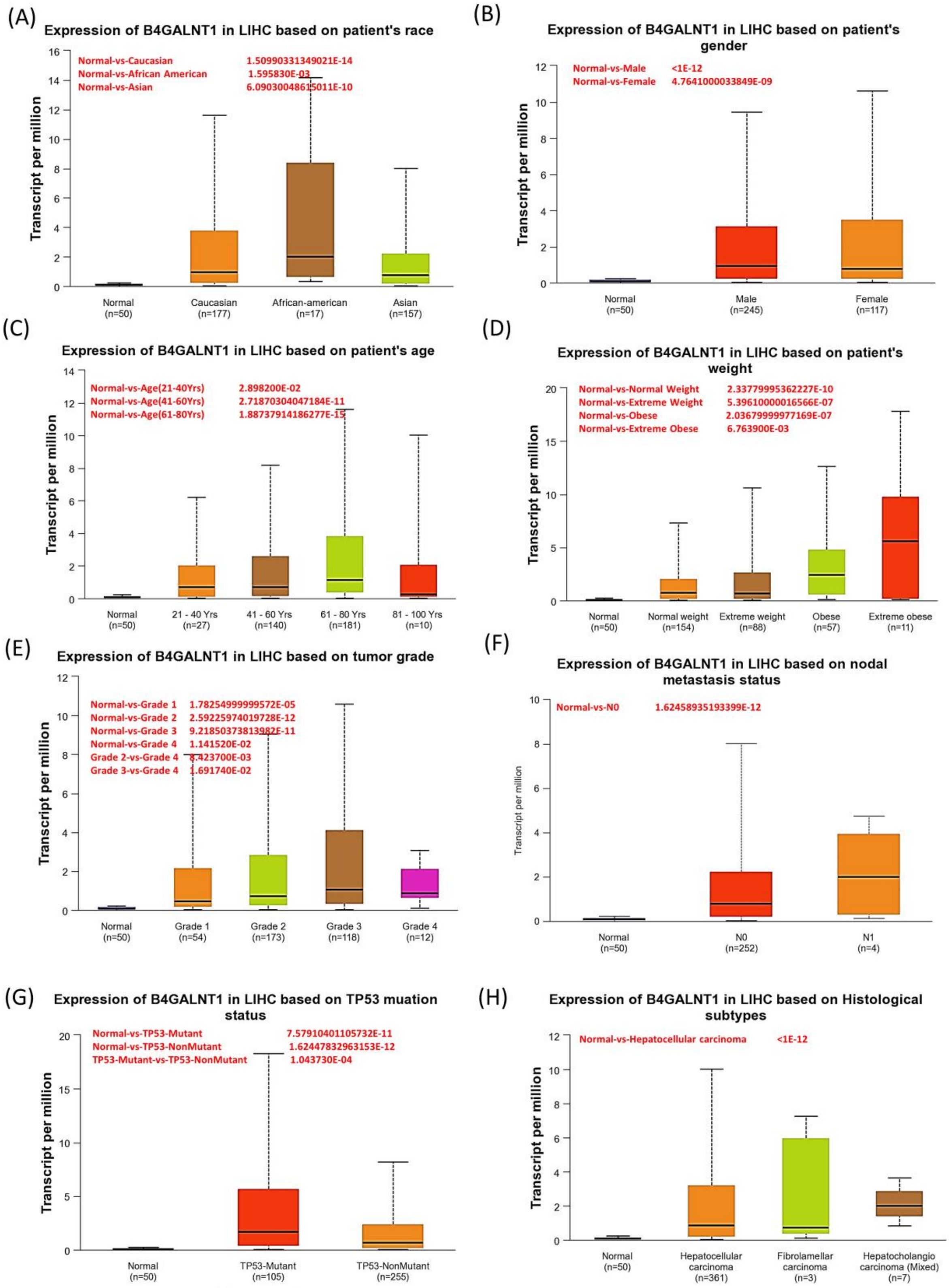


**Figure S1:** B4GALNT1 expression levels in LIHC cancers at various clinical stages (A) Race (B) Gender (C) Age (D) Weight (E) Tumor Grade (F) Nodal status (G) TP53 Mutation (H) Histological subtypes.

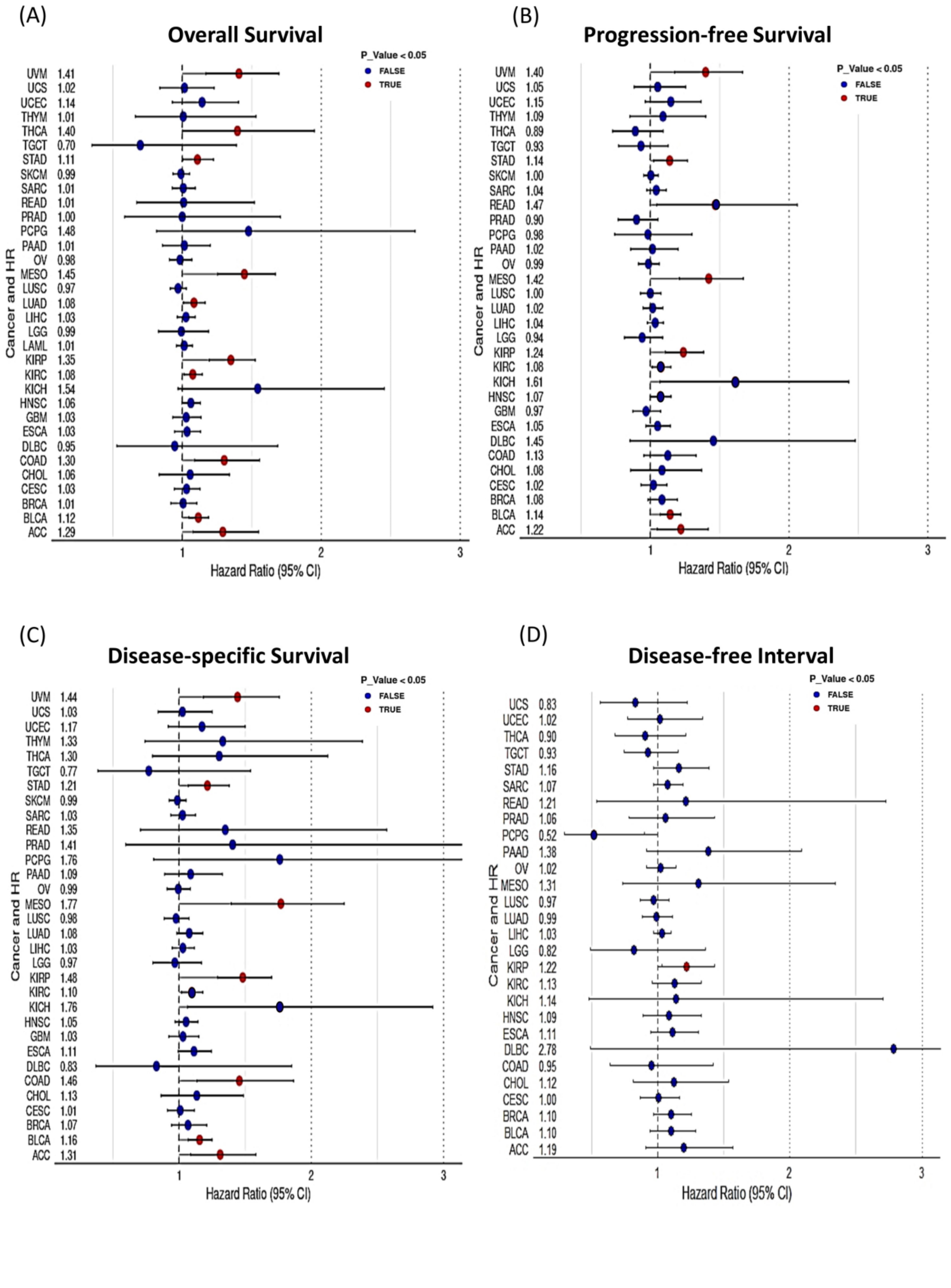


**Figure S2:** Survival analysis of B4GALNT1 in pan−cancer. (A) Overall survival (OS). (B) Progression−free survival (PFS) (C) Disease−specific survival (DSS). (D) Disease−free survival (DFS).

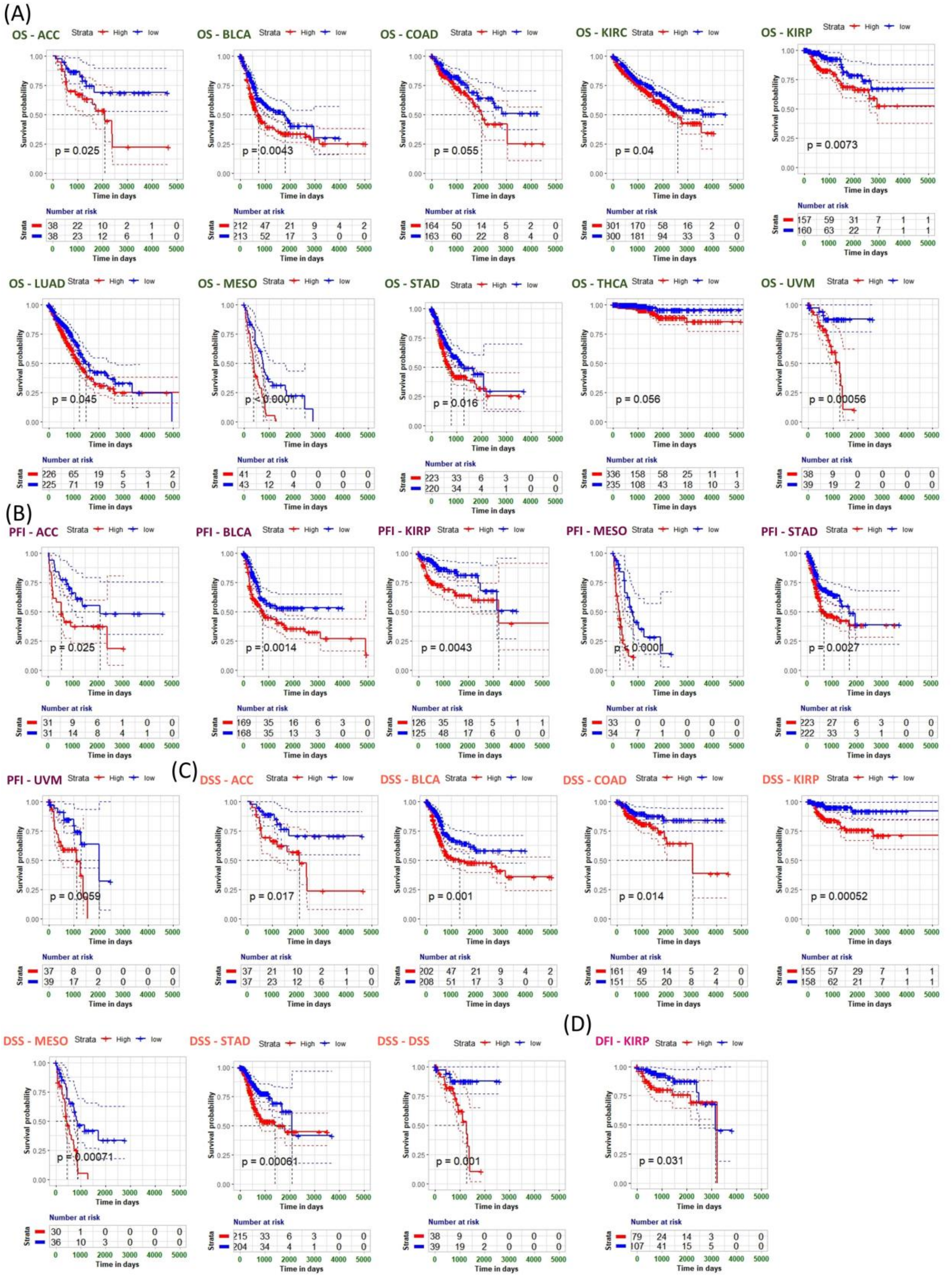


**Figure S3:** Association between B4GALNT1 expression and overall survival (OS). Progression−free survival (PFS) (C) Disease−specific survival (DSS). (D) Disease−free survival (DFS). Kaplan−Meier analysis of the association between B4GALNT1 expression with most significant differences in TCGA cancer ($p < 0.05$)

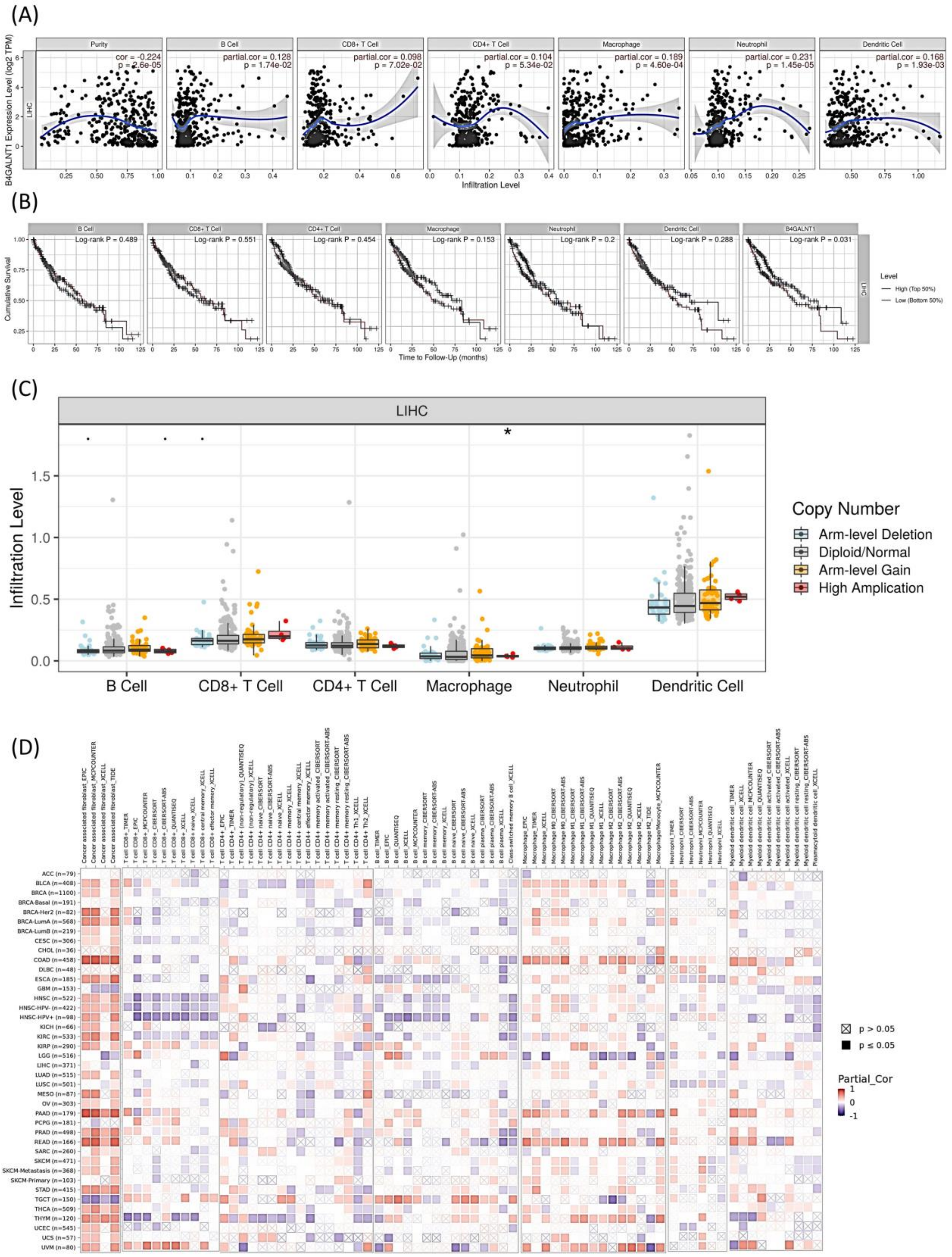


**Figure S4:** Correlation analysis with immune cells (A) Correlation of B4GALNT1 and immune infiltration of B cells, T cells CD8+, CD4+, Macrophages, Neutrophils and Dendritic cells in GI tract cancers, (B) Survival plot of immune infiltrates of B4GALNT1 expression, and (C) Somatic copy number alterations of B4GALNT1 in different TCGA cancers and their correlation of immune infiltrates.

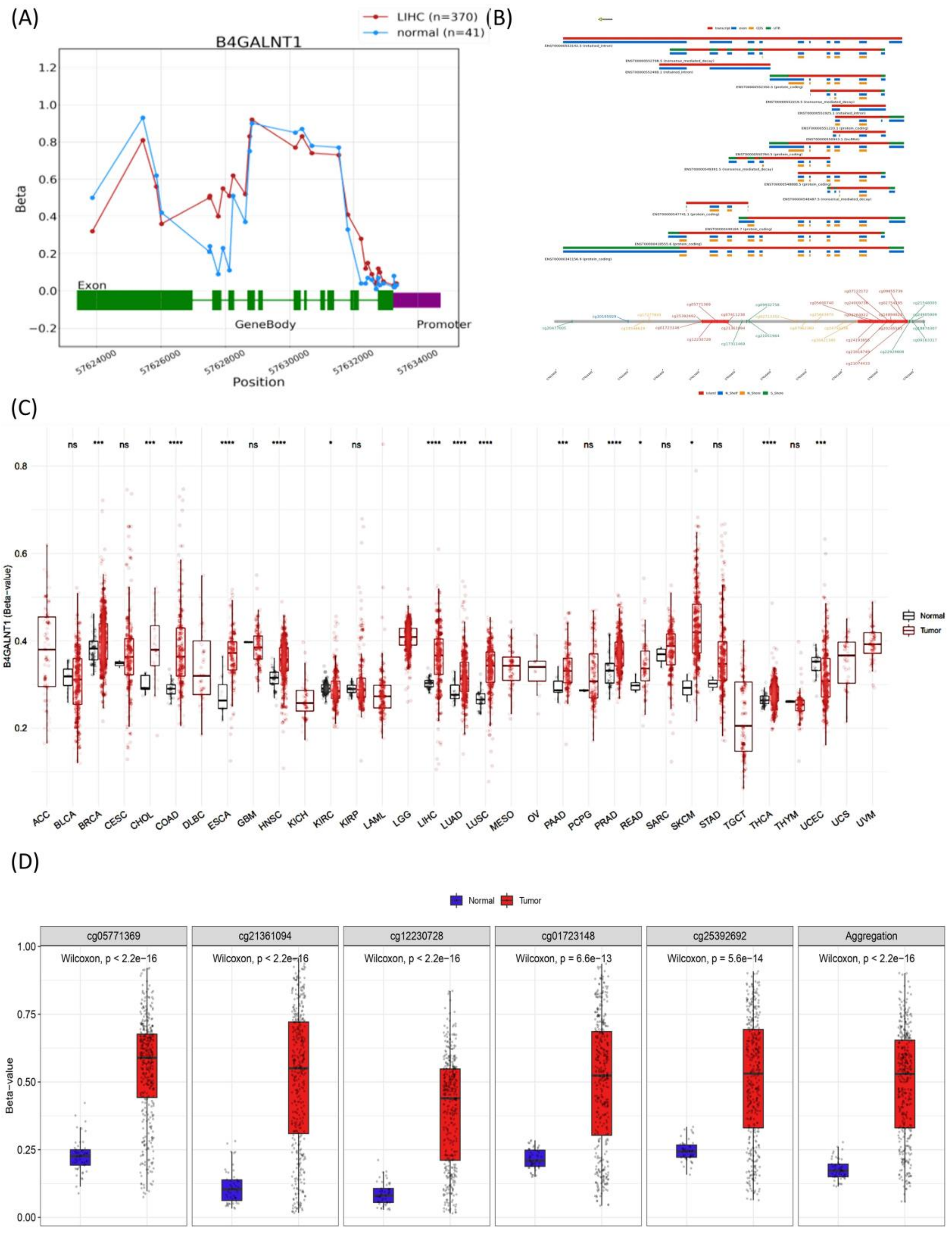


**Figure S5**: DNA methylation study of B4GALNT1 (A) Methylation in Promoter region, (B) Genomic information of B4GALNT1 the name of transcript is given, CpG shelves and shores, (C) CpG methylation aggegation of B4GALNT1 in tumor and normal samples, and (D) CpG probe expression in tumor and normal samples associated with B4GALNT1.

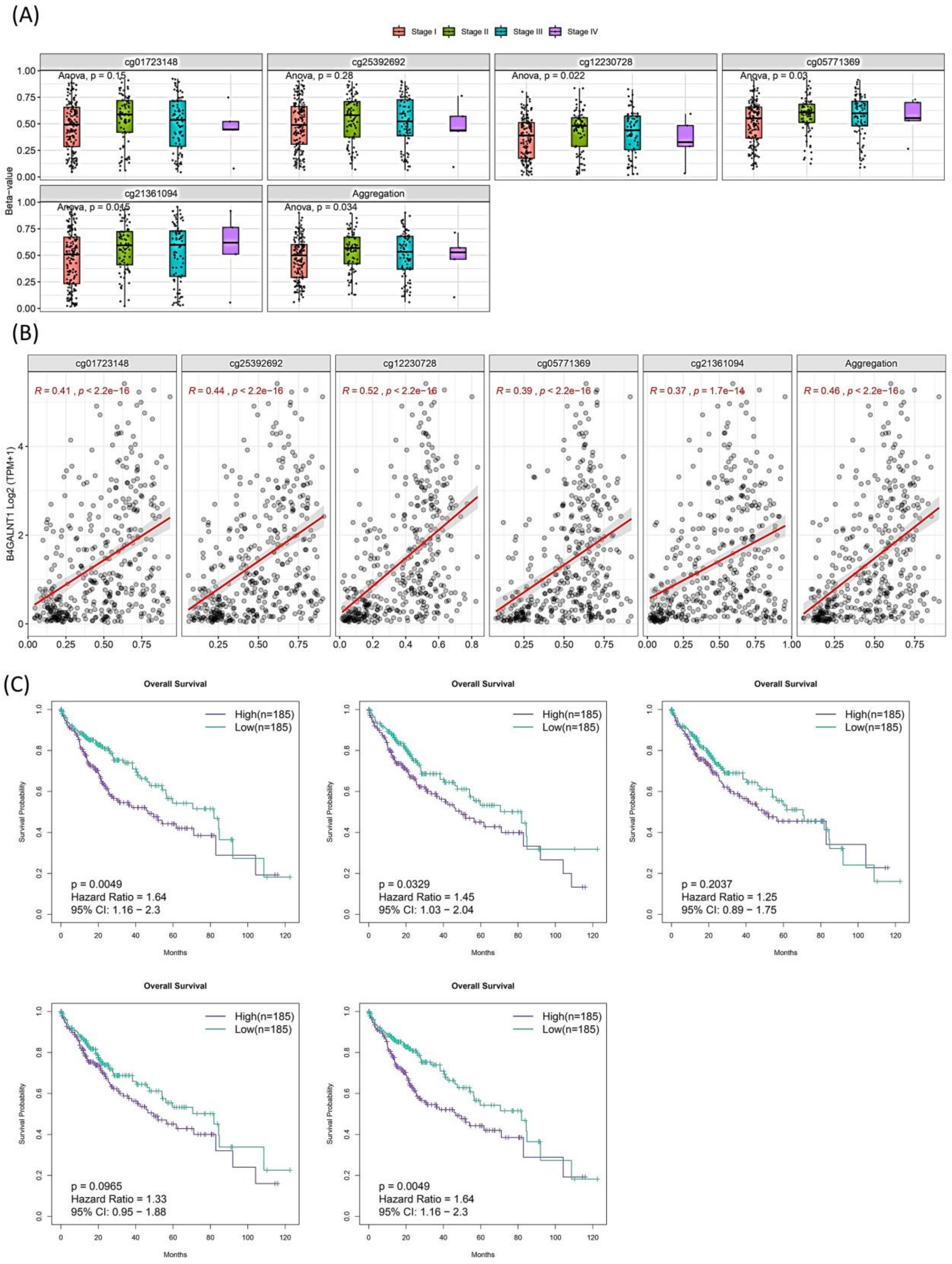


**Figure S6:** The methylation value compared across different stages for probe IDs (cg01723148, cg25392692, cg12230728, cg05771369 and cg21361094) , correlation with

B4GALNT1 and KM survival plot (A) Box plot comparison of different stages (B) Correlation plot of CpG probe Ids and B4GALNT1 expression (C) Survival plot of the CpG probe Ids.

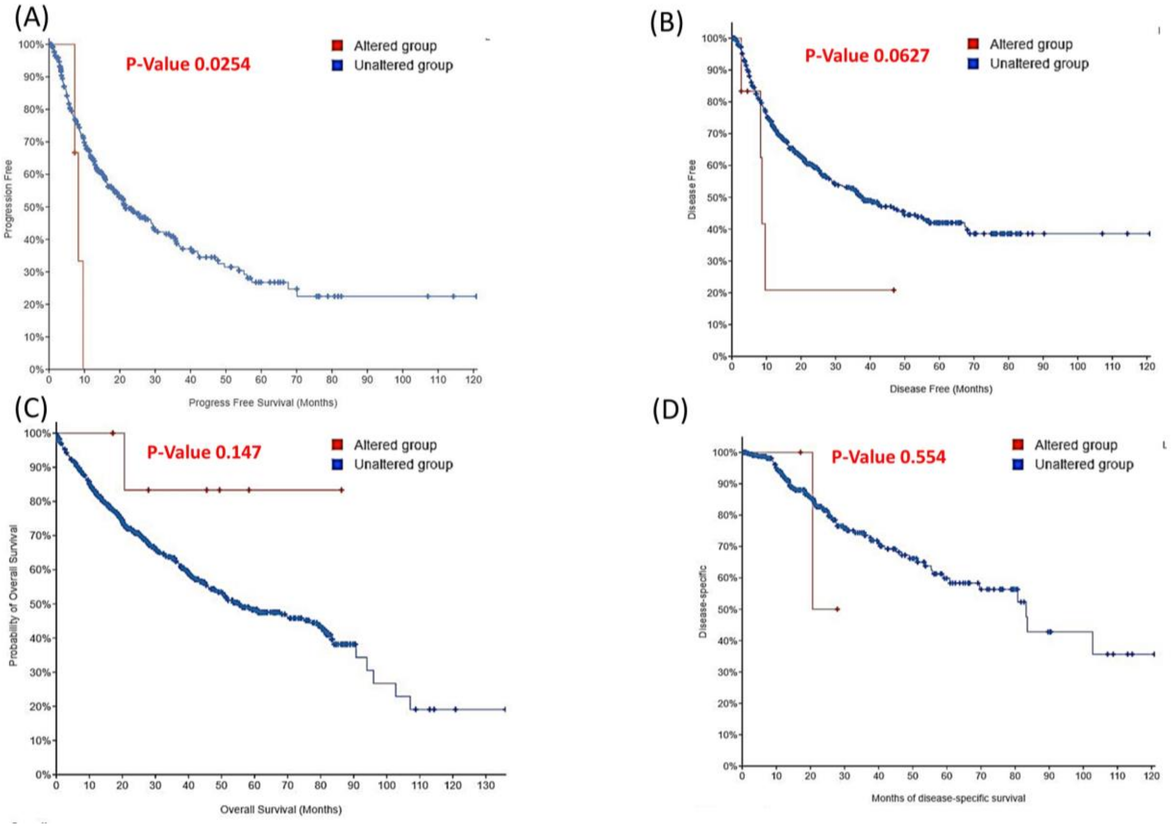


**Figure S7:** The survival analysis plot for altered and unaltered group for progression free, disease free, overall and disease specific analysis (A-D).
